# Origins and diversity of Exmoor ponies: A mitogenome framework for assessing maternal lineage diversification in endangered pony breeds

**DOI:** 10.1101/2025.10.30.685304

**Authors:** Deborah Davy, Anna Kinghorn, Alexandra Brumwell, Elizabeth Kilbride, Susan Baker, Philip M.C. Davy, Fatema Al-Ghaithi, Barbara K. Mable

**Affiliations:** School of Biodiversity, One Health & Veterinary Medicine, University of Glasgow, United Kingdom G20 6DE; Durrell Institute of Conservation and Ecology, School of Natural Sciences, University of Kent, United Kingdom CT2 7NR; Archivist for the Exmoor Pony Society; John A. Burns School of Medicine, University of Hawaii, USA; Faculty of Science, University of Porto, Portugal

## Abstract

In response to growing concerns over human-driven biodiversity loss, rewilding efforts increasingly employ semi-wild herbivores to restore ecosystems. The Exmoor pony is often cited as an ancient breed retaining primitive traits suited for such projects, yet its genetic history remains underexplored. We combined studbook records, mitochondrial DNA (mtDNA) D-loop, and whole mitogenome sequences to evaluate maternal lineage diversity for improved management recommendations and the phylogenetic placement of Exmoor ponies among modern and ancient breeds. We also assessed whether data from multiple breeds already available based on D-loop sequences could be organised into a useful framework for resolving the complex history of modern horse breeds. We identified eight mtDNA haplotypes among 88 Exmoor pony samples, representing all extant maternal lines. While phylogenetic trees based on D-loop sequences lacked resolution, clustering published sequences to a mitogenome-derived haplogroup framework allowed assessment of the distribution and relationships of lineages across diverse horse populations, including ancient DNA samples and Przelwalski’s horses. The most basal lineages of Exmoors were traced back to the Pleistocene, supporting their ancient origin and status as a primitive population, with a possible ancestral role in the development of some modern breeds. The founder lineages were found to be polyphyletic, with more derived haplogroups representing subsequent mixture with other domestic horse breeds. Although nuclear genome sequencing is needed to fully assess levels of admixture, we identified new haplotypes, along with rare or basal haplotypes that could be prioritised for conservation in rewilding studies. Our results highlight the potential for broad testing of hypotheses about origins and relationships among horse breeds using short regions of DNA mapped onto a more robust phylogenetic framework resolved through comparison of whole mitogenomes.

## Introduction

Current anthropogenically-driven biodiversity losses are characterized by increased extinction rates and a drastic loss of habitats (Díaz and Malhi 2022). In face of such alarming events, there has been growing interest in conservation approaches that emphasise active, flexible restoration of habitats like rewilding, which aims to restore ecosystems (Dobson et al. 1997; Carver 2016; Díaz and Malhi 2022) to a “self-sustaining state with natural ecological processes and minimal human intervention” (Perino et al. 2019). There have been several landmark projects, such as the Knepp Rewilding Project in the United Kingdom (UK), which has led to further popular interest and discussion of this practice (Lorimer et al. 2015; Sandom, Christopher J. and Wynne-Jones 2019). One of the key features of rewilding is the (re)introduction of species that restore ecosystem functioning (Sandom, Chris et al. 2013). Some resilient breeds of horses are particularly relevant for this practice as they are selective grazers that feed on abrasive grasses and herbs, resulting in mosaics of high and low vegetation and diverse habitats (Naundrup and Svenning 2015). Although guidelines for rewilding suggest inclusion of multiple lineages and genetic exchange through translocation of individuals between rewilding areas (Linnartz et al. 2023), these have so far mostly relied on pedigree records (when available) rather than genetic data.

However, previous studies have found horse studbooks to contain some erroneous pedigrees and, thus, be unrepresentative of the actual gene pool (Giontella et al. 2020). Consequently, establishing heritage of individuals used in conservation management is critical to avoid loss of rare variants or accidental breeding of related individuals. One breed that this could be important for is the Exmoor pony, a putatively ancient population native to the British Isles (Speed and Speed 1977; Baker 2008; Hovens and Rijkers 2013). They have been promoted for conservation grazing projects (Exmoor Pony Society 2025b; Rare Breeds Survival Trust 2025a) but without considering genetic variation. They have been naturally selected to survive the harsh conditions of the moorland of Exmoor (Gates 1979), and, due to their specialised grazing habits, they are crucial ecosystem engineers (Davy 2024). At present, they are one of the most endangered horse breeds, included as a priority species in the Rare Breed Survival Trust watchlist (Rare Breeds Survival Trust 2025b), with a population of approximately 500 individuals on Exmoor, and another 3500 individuals across the UK and various other countries (Exmoor Pony Society 2025c). However, they experienced a severe bottleneck starting at the end of WWII, with fewer than 50 individuals remaining, but not all contributed to future generations; consequently, based on genome-wide single nucleotide polymorphism (SNP) genotyping data, the Exmoor pony breed is highly inbred, with lower levels of genetic diversity compared to other breeds (Petersen et al. 2013). Based on studbook records (Exmoor Pony Society 2025a), the current Exmoor population is reported to have descended from 41 female founders and three male founders, with uneven representation of these founder lines; however, their relationships to one another have not been established. Therefore, to maintain the long-term genetic health of the population, it is important to understand the ancestry of individuals in relation to each other, to prioritise which lineages should be the focus of conservation grazing and ecological restoration initiatives, and to develop strategies for maintaining genetic variation through targeted translocations.

However, there could be a trade-off between preserving genetic variation and breeding out adaptively important traits in breeds or populations that have been genetically isolated (Mable 2019; Mellya et al. 2025). Exmoor ponies have been considered to be closely related to extinct relatives of the domestic horse (Speed and Speed 1977; Goodwin et al. 2008), suggesting they could have a distinct genetic history from other horse breeds. The retention of primitive features has been thought to be due to isolation of Exmoors from other horse breeds (Hovens and Rijkers 2013). However, so far there has been a lack of phylogenetic evidence to support this. Based on mitochondrial D-loop sequences (also known as the control region), McGahern et al. (2006) assigned 962 horses from 75 breeds to seven major haplogroups and found that the 20 Exmoor samples included were allocated to two different haplogroups, both of which were shared with several other breeds, suggesting shared ancestry. A subset of these Exmoor sequences (n=12) was taken from Jansen et al. (2002), who resolved 25 horse breeds into 17 haplotypes, with Exmoor ponies placed in a cluster with other northern European ponies like Fjord, Icelandic, and Scottish Highland. Winton et al. (2020) performed a detailed analysis of ancestry of other putatively ancient British pony breeds but did not include Exmoors. Importantly, no previous studies have included samples representative of all founding Exmoor herds to identify rare or distinctive maternal lineages that should be preserved.

In order to resolve the ancestry of Exmoors, it is also necessary to place them into the historical context of other horse breeds. The population dynamics of domestication are highly complicated, making phylogenetic history of domesticated species difficult to determine compared to wild species (Bruford et al. 2003; Frantz et al. 2020), but particularly so for the domestic horse (*Equus ferus caballus*). It was thought to be domesticated from its wild ancestor (*Equus ferus*) approximately 5000-6000 years ago in the Eurasian Steppes (Ludwig et al. 2009; Petersen et al. 2013); it is likely that multiple domestication events subsequently took place across Eurasia (Vilà et al. 2001; Cieslak et al. 2010). This means there was a continuation of gene flow between domestic and wild horses, leading to higher levels of genetic diversity than is usual for domestic animals (Petersen et al. 2013). However, selective breeding led to the expansion of domestic breeds and extinction of many wild breeds of horses (Librado et al. 2016). In addition, high levels of gene flow between domesticated lineages have led to small evolutionary distances between breeds (Lippold et al. 2011). This has led to most previous mtDNA studies into the phylogeny of horses producing trees with little or no statistical support, and little or no phylogenetic structure (Vilà et al. 2001; Jansen et al. 2002; Royo et al. 2005; Cieslak et al. 2010; Lippold et al. 2011; Winton et al. 2020), preventing the identification of evolutionary relationships between breeds.

A recent meta-analysis of population-level studies on horse evolutionary relationships suggested that this complex history means that previous haplogroup-based classification systems are inadequate to capture genetic diversity and origins (Agbani et al. 2025). However, this could be because many of the studies that have attempted comprehensive assessment of maternal genetic diversity across horse breeds (Hill et al. 2002; McGahern et al. 2006; Winton et al. 2020) have been based only on the mitochondrial control region (often referred to as the D-loop). The relatively few studies that have considered whole mitogenome sequences have found better resolution of phylogenetic trees (Achilli et al. 2012; Kvist and Niskanen 2020), but have tended to sample only a few individuals per breed or to report only unique haplotypes rather than frequencies of haplotypes within and between breeds. Combining a mitogenome-based framework with broader sampling of individual breeds based on smaller regions of mtDNA could help to resolve historical relationships compared to considering the D-loop alone.

Morphological studies into the origins of the domestic horse also have suggested a polyphyletic ancestry. One hypothesis concerning the origin of modern horses suggests that waves of migration of early horses across the Bering Straits land bridge led to them developing into four distinct primitive ancestral types, varying according to their geographical destination (Schafer 1981): A) Northern group horses, divided into the Universal Pony type (Type 1 - Primeval Pony), originally distributed from the Steppes to the Atlantic seaboard and the Ancestral cold blood (Type 2 - Heavy Horses), adapted to Central and Northern European conditions; and B) Southern group horses, divided into the Steppe horse (Type 3), adapted to Central Eurasian conditions and proto-Arab horses (Type 4), adapted to North African and Middle Eastern conditions. Schafer (1981) suggested that modern breeds were developed from mixtures of these ancestral types and that the modern Exmoor pony closely resembled the Universal Pony type and was likely its most direct descendant, giving support to the idea that Exmoors are an ancient, un-mixed breed. However, most molecular studies have not specifically considered these morphological classifications in their analyses, and so further study using molecular evidence could determine if these four ancestral types have monophyletic origins that could inform perspectives on origins of ancient lineages.

In reality, the evolutionary history of even highly morphologically distinctive ancient breeds remains unclear. For instance, Przewalski’s horse (*Equus ferus przelwalskii*; also referred to as Mongolian wild horse, Takhi) is native to the Eurasian Steppes and historically has been considered to be a wild relative of the domestic horse, with a distinct, ancient ancestry; however, this remains uncertain, given that it went extinct in the wild in the 1960s (Turghan et al. 2022). Some mtDNA studies have suggested Przewalski’s horse has a shared genetic history with the domestic horse, and that they are more closely related than traditionally thought (Ishida et al. 1995; Lau et al. 2009; Kvist and Niskanen 2020), whereas genome sequencing from ancient DNA suggests that admixture occurred after domestication (Orlando 2020). Although mtDNA is not informative about admixture, it can provide useful perspectives on diversity of maternal origins for rare or endangered lineages like Przeswalskii’s horses or Exmoor ponies. Advances in approaches for isolating “ancient DNA” from fossil or museum samples provides the opportunity to assess the relative persistence of various lineages over time, in order to assess hypotheses about origins of domestic and “wild” breeds (Cieslak et al. 2010; Vilstrup et al. 2013).

The overall aims of this study were to investigate whether genetic data could be used to inform management of Exmoor pony breeding strategies and to determine the phylogenetic history of the Exmoor pony in relation to other horse and pony breeds. Specifically, this study combined stud book and historic records with D-loop (control region) sequences from the mtDNA and whole mitochondrial genome sequencing (mitogenome) to: 1) verify matrilines based on observational pedigrees using mtDNA control region sequences; 2) quantify control region variation among Exmoor maternal lineages, in relation to a broader context of published sequences; 3) set the phylogenetic context of Exmoor pony maternal mitogenome haplotypes in relation to other breeds, using the framework proposed by Achilli et al. (2012); 4) determine whether D-loop sequences alone can be used to assign breeds to mitogenome haplogroups; 5) determine the relative frequency of haplogroups among horse breeds and breed groups to assess whether putatively ancient pony breeds such as Exmoors are differentiated from other horse breed groups; and 6) assess the relative persistence of haplogroups over time based on published D-loop sequences obtained from ancient samples. Although the study is focused on Exmoor ponies, the general approach used could have broad applicability across other domesticated breeds with complex evolutionary histories.

## Methods

### Pedigree Analysis

The Exmoor pony society was formed in 1921, when ponies that met the defined breed standard were registered in the studbook. Some of these had parents that had been registered pre-1921 (Polo Pony Stud Book 1894-1913, later renamed as the National Pony Stud Book) whereas others simply had herd of origin and were entered as “foundation stock” with unknown parents. The Exmoor Pony Study Book was closed in 1961; only ponies with sire and dam registered prior to that were eligible for inclusion. All existing individuals thus have descended from a limited number of founding herds: four main founder herds (in this study referred to as herds 900, 1, 12 and 23), as well as six other small herds (herds 27, 32, 44, 48, 54 and 76), thought to be subgroups (Baker 2008). Herds 27, 32, 44, 48 and 54 are subgroups of herd 23, and herd 76 has been hypothesised to be a subgroup of herd 1. Herd 1 and 900 are thought to have been closely connected during the 1950’s. The 2025 EPS studbook indicates that only 28 of the 41 maternal founder lines are still represented in the current population, although one line is from a founder accepted after inspection and an additional five lines are functionally extinct. To trace the founder lineage, matriline and patriline of Exmoor pony individuals, for this study, the EPS Stud Book records were accessed via The Exmoor Pony Society Stud Book, 3rd Edition, 1980, and the online Stud Book (https://breeds.grassroots.co.uk/Home), The National Pony Society Stud Book 1913-47, and breeders’ personal records.

Most ponies were identified by a herd number, a unique number allocated to their breeder by the EPS to identify all the ponies born into that herd, followed by that pony’s unique individual number within that herd; e.g., the founder ID 1020 indicates that the pony originated in Herd 1, and that her individual identity within the herd was 20 (Supplementary Table 1). However, ponies from the Anchor herd, where an anchor symbol is used instead of a herd identification number, were recorded in the Stud Book as 000. For clarity, we allocated them the herd number 900; therefore, the founder pony 000041, was individual 41 from the Anchor herd; her sampled descendent from the current Anchor herd has sample ID 900600 and is therefore Anchor herd pony, individual 600. The founder matriline IDs correspond to the herds of origin of the founder mares. The pony sampled ID and founder matriline ID will often be for different herds as they are multiple generations apart. The founder herds sold stock to other breeders who registered the progeny into their own herds, and foundation Herd 1 was closed in 1960. Founder representation was calculated using EPS Stud Book records, to show the number of live females and live potentially breeding males whose pedigrees can be traced to specific founders. Males known to be castrates or not approved for breeding were excluded. This information was then used in the selection of ponies for DNA sequencing; sampling prioritised coverage of as many founding matrilines as possible.

### Sampling

To obtain a comprehensive overview of the genetic diversity within the current Exmoor pony population in the UK, hair samples were obtained from all matrilineal founder lines represented in the post-WW2 founder population present in the UK, and from males representing the founder patrilines. When possible, samples were collected from multiple individuals representing each matrilineal founder line to improve the accuracy of the verification of the maternal haplotypes present (Table 1).

**Table 1.** Summary of dam lines sequenced, indicating the number of samples from each founder line (after correcting metadata errors), the D-loop haplotype frequency and the number of samples that also had full mitogenomes sequenced.

|  |  | D-Loop Haplotype |  |  |  |  |  |  |  |  |
| --- | --- | --- | --- | --- | --- | --- | --- | --- | --- | --- |
| Dam Line | N Samples | 1 | 2 | 3 | 4 | 5 | 6 | 7 | 8 | N Mitogenomes |
| 01/020 | 12 | 0 | 0 | 0 | 0 | 0 | 0 | 12 | 0 | 5 |
| 01/028 | 2 | 0 | 0 | 0 | 0 | 2 | 0 | 0 | 0 | 1 |
| 01/030 | 3 | 0 | 0 | 0 | 0 | 0 | 0 | 3 | 0 | 2 |
| 01/031 | 2 | 0 | 0 | 0 | 0 | 0 | 0 | 2 | 0 | 2 |
| 01/035 | 4 | 0 | 0 | 0 | 0 | 0 | 0 | 4 | 0 | 2 |
| 12/002 | 11 | 0 | 0 | 0 | 0 | 0 | 11 | 0 | 0 | 7 |
| 12/011 | 6 | 6 | 0 | 0 | 0 | 0 | 0 | 0 | 0 | 5 |
| 23/001 | 2 | 0 | 0 | 0 | 0 | 2 | 0 | 0 | 0 | 1 |
| 23/008 | 2 | 0 | 0 | 0 | 2 | 0 | 0 | 0 | 0 | 2 |
| 23/010 | 2 | 0 | 0 | 0 | 2 | 0 | 0 | 0 | 0 | 1 |
| 27/001 | 1 | 1 | 0 | 0 | 0 | 0 | 0 | 0 | 0 | 1 |
| 44/002 | 1 | 0 | 0 | 1 | 0 | 0 | 0 | 0 | 0 | 1 |
| 48/029 | 3 | 0 | 0 | 0 | 0 | 0 | 0 | 3 | 0 | 0 |
| 76/001 | 9 | 0 | 9 | 0 | 0 | 0 | 0 | 0 | 0 | 7 |
| 76/003 | 1 | 0 | 0 | 0 | 0 | 0 | 0 | 0 | 1 | 1 |
| 900/002 | 2 | 0 | 0 | 0 | 0 | 0 | 0 | 2 | 0 | 1 |
| 900/003 | 1 | 0 | 0 | 0 | 0 | 0 | 0 | 1 | 0 | 1 |
| 900/005 | 3 | 0 | 0 | 0 | 0 | 0 | 0 | 3 | 0 | 1 |
| 900/041 | 2 | 0 | 0 | 0 | 0 | 0 | 0 | 2 | 0 | 1 |
| 900/049 | 2 | 0 | 0 | 0 | 0 | 0 | 0 | 2 | 0 | 1 |
| 900/050 | 4 | 0 | 0 | 0 | 0 | 0 | 0 | 4 | 0 | 1 |
| 900/051 | 5 | 0 | 0 | 0 | 0 | 0 | 0 | 5 | 0 | 5 |
| 900/052 | 3 | 0 | 0 | 0 | 0 | 0 | 0 | 3 | 0 | 2 |
| 900/054 | 5 | 1 | 0 | 0 | 0 | 0 | 0 | 4 | 0 | 1 |
| Frequency | 88 | 8 | 9 | 1 | 4 | 4 | 11 | 50 | 1 | 52 |

Samples were collected by plucking 10-20 hairs with follicles attached from the tails or manes of the sampled individuals, the sampler changing gloves or cleaning hands between each collection to avoid cross-contamination. Samples were stored in labelled paper envelopes or sealed plastic bags until extraction of DNA from the hair follicles. Samples were collected from registered Exmoor ponies in various locations and herds from Scotland, England and Wales in the United Kingdom; the locations of individuals are not provided, to protect the privacy of the owners. Samples were obtained from 73 living ponies, along with archived samples from 15 dead individuals; this included a total of 67 females and 21 males, from ponies born between 1978 and 2021 (Supplementary Table 1).

### DNA Extractions

DNA was extracted using Qiagen DNeasy kits (Qiagen Inc, Paisley, UK), using the manufacturer’s protocols for hair samples. For each individual, 1 cm of hair was cut from the end of each hair in the sample with spring bow scissors. The hair was then transferred into a 1.5 ul Eppendorf tube containing 180 ul ATL (lysis) buffer, to digest the cells and release DNA, and 20 ul Proteinase K solution, to denature released proteins that may degrade the DNA. The hair was then chopped into smaller pieces and incubated at 56°C with shaking overnight. Between preparing each sample, the instruments used were treated with UV light to prevent contamination.

Each sample was treated with 4 ul RNase (100 mg/ul) and incubated at 37°C for 30 minutes to remove RNA. Samples were eluted from the spin columns in 42 ul AE elution buffer. To increase yields, this was repeated with 42 ul and then 22 ul elution buffer. For each set of extractions, a negative extraction control was included, to check for contamination. The concentration and purity of the DNA extractions were checked using a nanodrop ND-1000 spectrophotometer (Thermo-Fisher, Cambridge UK) and agarose gel electrophoresis. For samples used for whole genome sequencing, DNA was further quantified using a Qubit 2.0 Fluorometer (Thermo-Fisher, Cambridge UK).

### Mitochondrial D-loop Amplification and Sequencing

A 426-bp fragment of the mitochondrial control region was amplified using primers described in Hill et al. (2002): IRD700 5ʹ-CTA GCT CCA CCA TCA ACA CC-3ʹ and IRD800 5ʹ-ATG GCC CTG AAG AAA GAA CC-3ʹ. PCR was performed with a 20 ul master mix: 15.4 ul DNase free water, 2 ul 10x PCR buffer, 0.6 ul Mg (50 mM), 0.4 ul dNTPs (10 mM), 0.2 ul IRD700 primer (10 mM), 0.2 ul IRD 800 primer (10 mM), 0.2 ul Invitrogen *Taq* polymerase (5 Units/ul) and 1 ul DNA. Thermal cycling was carried out at 94°C for 5 minutes, followed by 30 cycles of 94°C for 60s, 50°C for 60s and 72°C for 60s, followed by a final extension at 72°C for 6 minutes. The negative extraction controls and a negative PCR control were included with each set of amplifications, to check for contamination. Amplified products were visualised using 2% agarose gel electrophoresis. PCR products were sent to the University of Dundee for Sanger sequencing service on an ABI 3750 DNA analyser, using forward and reverse primers for each sample.

### Mitogenome Sequencing

DNA extracted from a subset of 55 individuals was selected for whole genome sequencing (Table 1); these samples represented all paternal and all except one of the female founder lines used for D-loop sequencing (Supplementary Table 1); the DNA from one maternal lineage (48/029) was not obtained until after the whole genome sequencing had been completed. Samples were sent to Novogene UK and sequenced in three batches between 2020 and 2022 (Supplementary Table 2). The samples were sequenced on a Novaseq 6000 with 150 bp paired end chemistry, aiming for 27 Gb of raw data per sample, to achieve 10x coverage per individual.

The full code for the following pipelines can be found at https://github.com/alexbrumwell/Honours_project. Prior to data processing, read quality was assessed using FastQC v. 0.11.8 (Babraham Institute, 2018). Firstly, Trim Galore (https://www.bioinformatics.babraham.ac.uk/projects/trim_galore/) was used to trim the adaptors of the sequences and then the data was mapped to a horse mitogenome reference (NCBI reference: NC_001640.1) using BWA MEM v. 0.7.15 (Li and Durbin 2009). Next, SAMTOOLS v. 1.9 (Li et al. 2009) was used to convert the SAM files into BAM files and sort them and remove any reads that had a quality score under 30. SAMTOOLs fixmate and markdup were used to remove duplicate reads. SAMTOOLs merge was used to merge files that contained reads from the same individual. PICARDTOOLS v. 2.8.2 (Broad Institute) with the ValidateSamFile option was used to assess the validity of the BAMs. QUALIMAP bamqc (García-Alcalde et al. 2012) was used to calculate the average coverage of each individual. Then, ANGSD v. 0.916 (Korneliussen et al. 2014) was used with the -doFasta option to call the consensus FASTA for each individual, using the average coverage of each genome as the minimum depth.

### D-Loop Haplotype Assignment

The new Exmoor D-loop sequences (n = 88) were aligned and manually corrected using Sequencher 5.4.6 (Gene Codes Inc, Ann Arbor, Michigan). A consensus sequence was created for each individual sample where a forward and reverse sequence was present. Consensus sequences were then organized into unique haplotypes, using the “assemble automatically” function in Sequencher and setting the assembly parameters at minimum overlap 20 bp and 100% similarity. To visualize the distribution of haplotypes across maternal founding lineages (as recorded in the EPS studbook), the unique set of haplotypes and their frequencies was used to create a minimum spanning network using Popart (Bandelt et al. 1999).

To set the Exmoor maternal haplotypes identified into a broader context, the published mtDNA control region sequences of all available Exmoor ponies were downloaded from GenBank (n=32), along with 1548 other horse sequences consisting of 87 other domestic horse breeds, including 25 Przewalskii’s horse sequences. Although there have been over 70 papers that have included D-loop sequences of horses, the majority of sequences were downloaded by searching the nucleotide database of NCBI Entrez interface for the accession numbers from three previous mtDNA studies (Supplementary Table 3), which spanned a wide variety of different breeds from across Europe, Asia and Africa and included both new and previously published sequences: McGahern et al. (2006), Achilli et al. (2012), and Kvist and Niskanen (2020). The McGahern et al. (2006) set was comprehensive at the time for all available D-loop sequences from a wide range of global breeds (n = 962 sequences), including 20 Exmoors; Achilli et al. (2012) established a framework for mtDNA haplogroup resolution based on whole mitogenome sequences (n = 83 unique haplotypes; including only a single Exmoor sequence); Kvist and Niskanen (2020) listed additional mitogenome sequences, including 17 from Przewalskii’s horses (Der Sarkissian et al. 2015). In addition, NCBI BLAST was used to search for other matching sequences to the Exmoor haplotypes on Genbank, as well as searching NCBI nucleotide with the keywords “Exmoor” and “D-loop” or “control region”.

Horse breeds were classified according to the Schafer (1981) scheme: Type 2 - Heavy or Cold-Blood; Type 3 - Steppe; and Type 4 - Proto-Arab. We considered ponies as a separate category (i.e. “Type 1 - Primitive Pony”) from the horse breeds because most would have been classified as primitive (Supplementary Table 3).

Sequences from GenBank were added to the Sequencher file and aligned with the Exmoor consensus sequences using the “assemble automatically” function set with an overlap of at least 20 bp and 80% similarity. Since the published studies had not all sequenced the same part of the mtDNA control region, sequences which did not overlap the region sequenced for the Exmoors were removed from the alignment. Aligned sequences were visualized in MacClade 4.05 (Maddison and Maddison 2000) and manually inspected for errors and length variation. Although the Exmoor sequences were 426 bp in length, the final alignment was pruned to 350 bp, to avoid regions with ambiguities in published sequences but to include the region where the most published sequences overlapped. The pruned sequences were collapsed into unique haplotypes using DNAsp6.12.03 (Rozas et al. 2017) and the haplotype that each sequence was assigned to, along with the breed it had been assigned in Genbank was recorded. Singletons were checked for zero length branches based on a neighbor joining tree assembled in Mega 11 (Tamura et al. 2021) and added to haplotypes accordingly.

### Data Analysis

#### D-Loop Exmoor maternal lineage distribution and sharing with other breeds

To visualize the distribution of haplotypes across maternal founding lineages (as recorded in the EPS studbook), the unique set of Exmoor haplotypes and their frequencies was used to create a minimum spanning network using Popart (Bandelt et al. 1999). To determine the uniqueness of Exmoor haplotypes, the full set of sequences downloaded from Genbank were aligned to the Exmoor haplotypes and the frequency of sharing with various breeds was recorded.

#### Phylogenetic relationships based on mitogenomes

In order to set the context of evolutionary relationships among maternal lineages of Exmoor ponies, the mitogenomes of the 55 Exmoor pony whole genome sequences assembled here were aligned to the mitogenome sequences provided by Achilli et al. (2012), along with the *Equus caballus* reference mitochondrial genome (NC_001640.1), using Aliview version 1.28 (Larsson 2014). Achilli et al. (2012) had included only a single Exmoor sequence and no other Exmoor mitogenomes were available through Genbank. Additional published mitogenome sequences from other breeds were then added from a list provided by Kvist and Niskanen (2020), which included sequences from Przelwalski’s horses (Der Sarkissian et al. 2015), a thoroughbred and a Yakut breed. All of these had also been included in the analysis of D-loop haplotype variation above. Sequences were manually corrected for indel errors (mostly in homopolymers) and collapsed into unique haplotypes using DNAsp6.12.03 (Rozas et al. 2017). Haplotypes that differed from one another only within a microsatellite repeat region in the D-loop (between bp 16124 and 16354) were combined with the consensus haplotype for the rest of the mitogenome. The best-fitting model of evolution of the final set of unique haplotypes was determined using model test, as implemented in Mega 11 (Tamura et al. 2021), excluding the microsatellite repeat region and using pairwise deletions for indels or missing data. A maximum likelihood tree with 1000 bootstrap pseudo-replications was generated using the parameters determined by model test, and pairwise deletions.

The tree was exported in Newick format and visualised using Evolview v2 (He et al. 2016) to determine whether the Exmoor mitogenomes and those taken from Qvist and Niskanen (2020) could be clearly mapped onto the haplogroups defined by Achilli et al. (2012), based on reciprocally monophyletic groups resolved with bootstrap support of at least 70%.

#### Phylogenetic relationships based on D-loop sequences

A phylogenetic tree was also constructed based on unique D-loop haplotype sequences, to determine whether clustering into the framework predicted by the Achilli et al. (2012) mitogenome was possible based on this short region of mtDNA alone. The best-fitting model of evolution was determined as for the mitogenome sequences and a minimum evolution (ME) tree with 1000 bootstrap replications was constructed, using the parameters estimated.

#### Distribution of mitogenome haplogroups across horse breeds

In order to assess the relative contribution of different maternal haplogroups to individual breeds (including Exmoors), each of the D loop haplotypes was assigned to a mitogenome haplogroup, if they fell into the reciprocally monophyletic groups on the D-loop phylogeny. The frequency of individual sequences falling into each haplogroup was then calculated. A phylogenetic tree generated from a single (arbitrarily selected) sequence from each of the Achilli et al. (2012) haplogroups was constructed in MEGA, to provide a framework for visualising relative frequencies in various breeds and breed groups (i.e. based on the morphological classifications from Schafer 1981), using frequency heat maps mapped onto the tree in Evolview v2 (He et al. 2016).

“Orphan” singletons that could not be unambiguously assigned to a haplotype due to missing data were excluded from the D-loop tree but were assigned to haplogroups based on clustering in a minimum evolution tree (Supplementary Table 3). Winton et al. (2020) sequenced the D-loop for 484 British ponies (but excluding Exmoors) but their sequence set was missing the first 37 bp of our alignment, which included some key polymorphic sites to distinguish individual haplotypes. These were thus included in calculations of relative frequency of sequences matching Exmoor haplotypes and mitogenome haplogroup frequencies but not in the phylogenetic tree.

#### Persistence of haplogroups over time

To assess the relative persistence of haplogroups over time, we also downloaded and aligned D-loop sequences from “ancient DNA” samples newly sequenced or compiled by Cieslak et al. (2010) from a set of previous studies (Di Bernardo et al. 2004; Keyser-Tracqui et al. 2005; Weinstock et al. 2005; McGahern et al. 2006; Cai et al. 2007; Lira et al. 2010; Priskin et al. 2010). These were classified in relation to age and broad geographic regions rather than breeds (Supplementary Table 4). Sequences were assigned to mitogenome haplogroups using minimum evolution-based clustering, as for the modern samples. A backbone phylogeny was created (using maximum likelihood with 500 bootstrap pseudo-replications) using the D-loop region of the mitogenome haplogroup representatives, with the addition of any sequences that were unique to the ancient samples, to visualise the relative frequency of haplogroups in different eras.

## Results

### D-Loop Exmoor maternal lineage distribution and sharing with other breeds

A total of eight unique haplotypes were found among the 88 Exmoor individuals sequenced, based on 350 bp of D-loop (control region) mtDNA sequences (Table 1). There were 32 polymorphic sites, with the most similar haplotypes differing by 2 bp (haplotypes 1 and 2) and the most divergent differing by 14 bp (haplotypes 6 and 8) (Supplementary Table 5). Average pairwise nucleotide diversity (pi) across the eight haplotypes was 0.028. The samples spanned 24 founder matrilines, with haplotype 7 being found at the highest frequency and shared by 14 of the lineages (Figure 1; Table 1). All other haplotypes were restricted to 1-3 maternal lineages. Five maternal founding lineages (1020, 1035, 12002, 12011, and 900054) originally appeared to have two haplotypes, which should not be possible for mtDNA (Supplementary Figure 1). In each case, additional archive consultation revealed errors in the metadata associated with the hair samples. After correcting, all samples matched predictions about founding maternal lineages from the studbook records.

**Figure 1.**
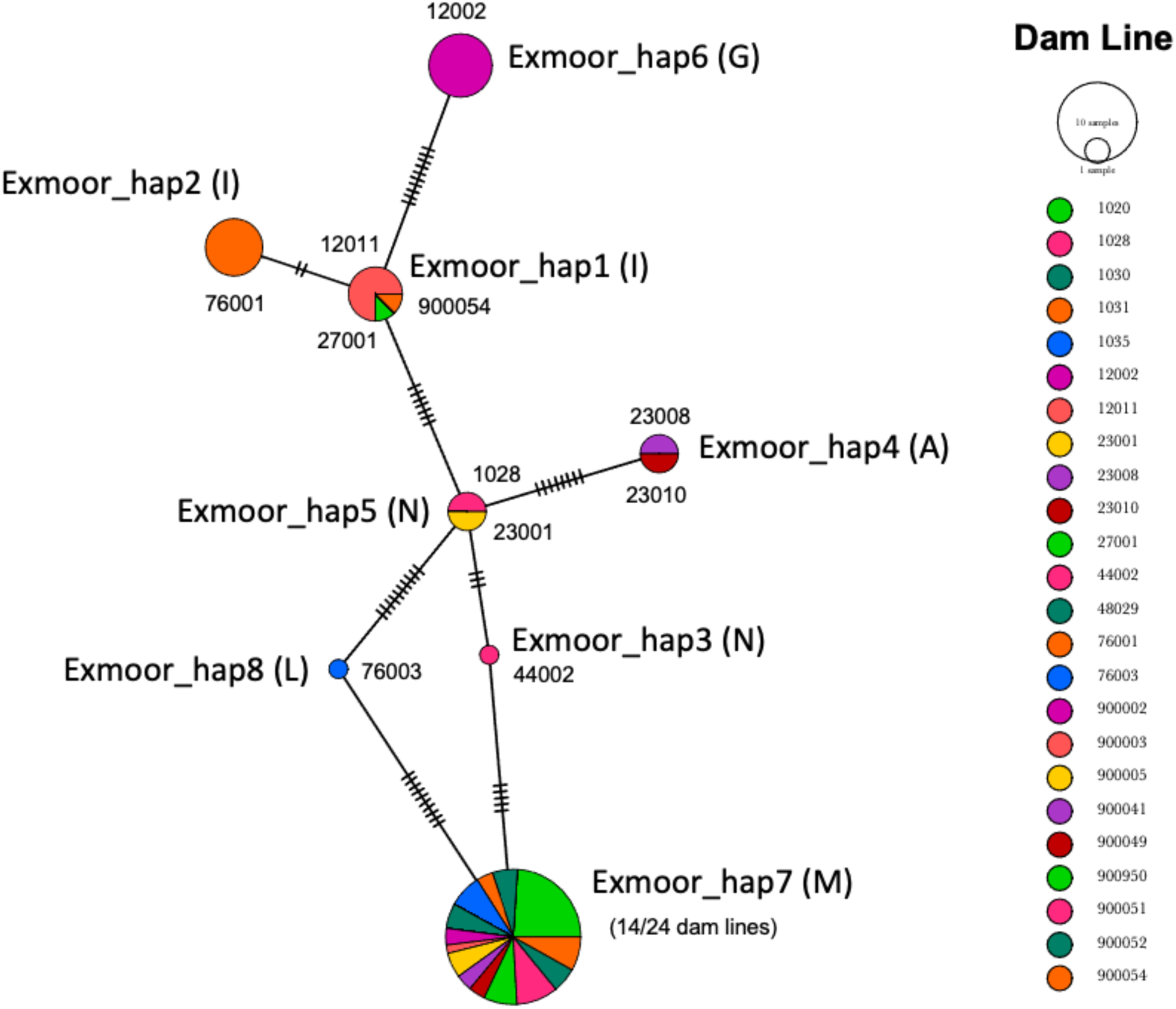
Minimum spanning D-loop haplotype network of newly sequenced Exmoor pony sequences, grouped by maternal founder lines (Dam lines). The eight Exmoor mitogenome haplotypes are indicated, along with the haplogroup (in parentheses) to which they were assigned (Achilli et al.2012). Size of the circles is proportional to the sample size; colours indicate maternal lineages; hatch marks indicate the number of mutations separating each haplotype. Based on the number of connections to other lineages, haplotype 5 (haplogroup N, found in two newly sequenced Exmoors from each of maternal lineages1028 and23001, three published Exmoors, and shared with other breeds: see Supplementary Table2), would be predicted to be ancestral. However, haplotype 7 was shared across more Exmoor maternal lineages (14/24 dam lines) and was shared with non-Exmoor breeds. Exmoor haplotypes 1 and 6 were not shared with other breeds and haplotype 2 was only shared with Welsh ponies. Haplotype 8 was only found in a single newly sequenced Exmoor but was shared with a wide range of other breeds (see Table2).

Based on the number of connecting haplotypes, haplotype 5 would be considered as ancestral in the haplotype network (Figure 1) but haplotype 7 was shared by the most lineages and was found at highest frequency. Haplotypes 1, 2 and 6 formed a distinctive cluster from the other haplotypes. Haplotype 8 also was highly divergent, differing from its most similar haplotypes (5 and 7) by 10 bp.

Two additional Exmoor haplotypes (Exmoor_hap9; Exmoor_hap10) were identified among published sequences but these were singletons, not shared with other breeds or any of the newly sequenced Exmoors (Supplementary Tables 3 and 5). All other published Exmoor sequences were identical to one of the eight newly sequenced Exmoor haplotypes.

Exmoor haplotypes 1 and 6 were only found in Exmoor samples; all other Exmoor haplotypes were shared with other breeds (Table 2; Supplementary Table 5). Exmoor_hap2 was shared with Welsh pony samples but from the Winton et al. (2020) study, which excluded the first 37 bp of the D-loop; thus, these could also be hap14, which was found in Przelswalskii’s horses. Exmoor_hap 4 was found at low frequency in other pony breeds (Scottish Highland and Welsh) and Exmoor_hap3 was found at low frequency in two heavy horse breeds (Merens and Orlov). Exmoor_haps7 and 8 were found across a broader range of breeds, spanning all breed groups.

**Table 2.**
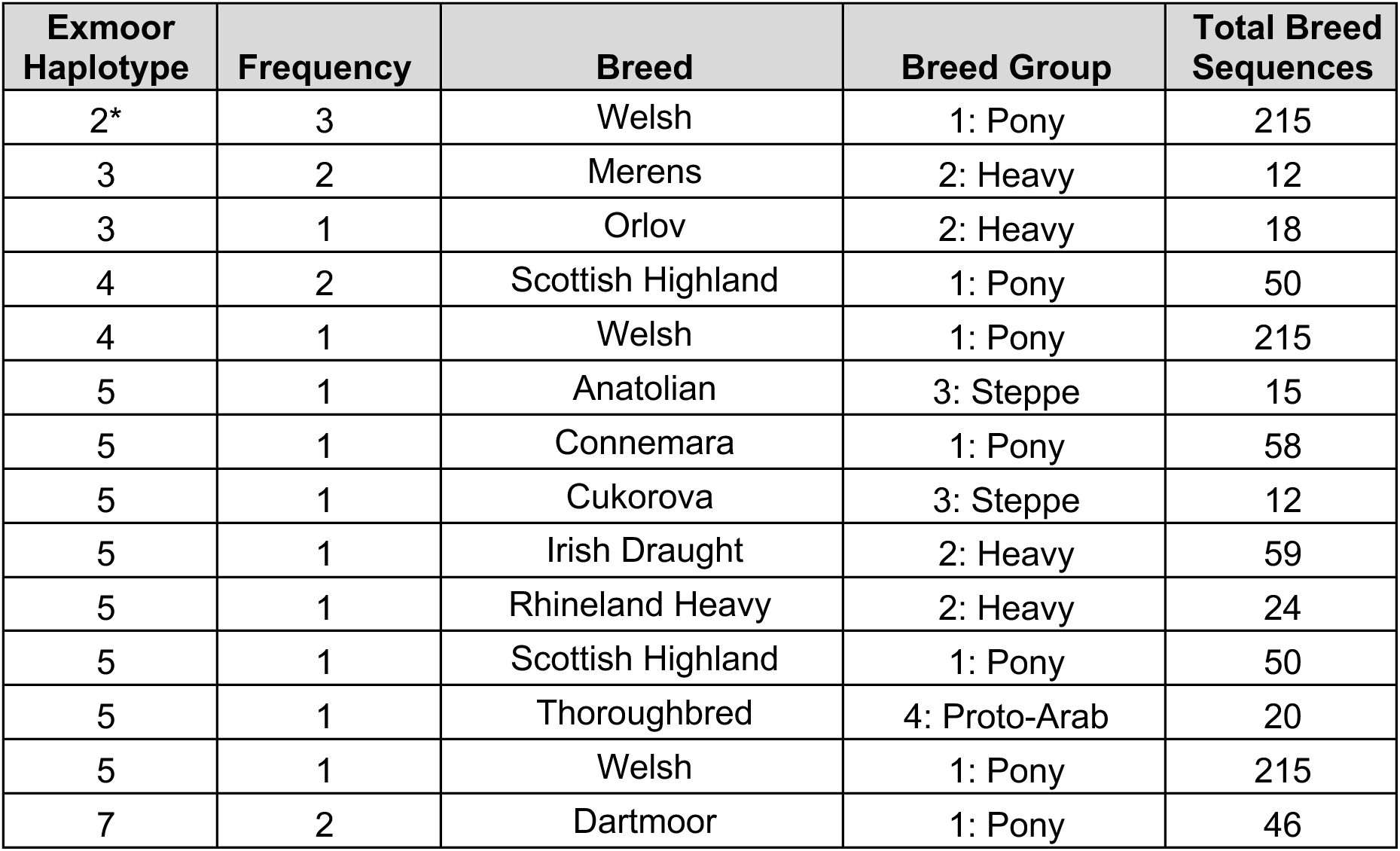
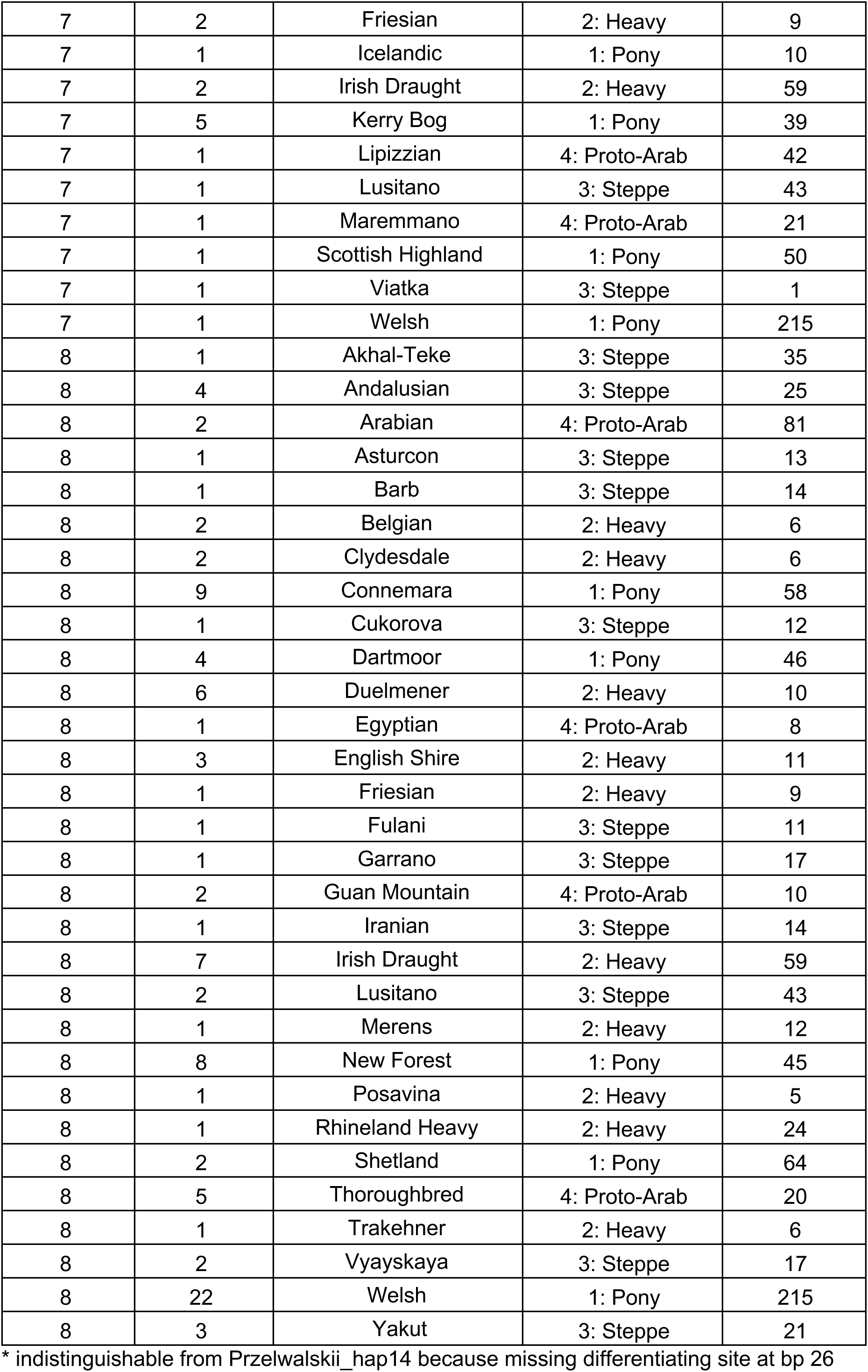
Haplotype sharing of Exmoor haplotypes with other breeds and breed groups, showing the frequency of the haplotype in the other breeds, their breed groups, and the total number of sequences that had been included from that breed.

| Exmoor Haplotype | Frequency | Breed | Breed Group | Total Breed Sequences |
| --- | --- | --- | --- | --- |
| 2* | 3 | Welsh | 1: Pony | 215 |
| 3 | 2 | Merens | 2: Heavy | 12 |
| 3 | 1 | Orlov | 2: Heavy | 18 |
| 4 | 2 | Scottish Highland | 1: Pony | 50 |
| 4 | 1 | Welsh | 1: Pony | 215 |
| 5 | 1 | Anatolian | 3: Steppe | 15 |
| 5 | 1 | Connemara | 1: Pony | 58 |
| 5 | 1 | Cukorova | 3: Steppe | 12 |
| 5 | 1 | Irish Draught | 2: Heavy | 59 |
| 5 | 1 | Rhineland Heavy | 2: Heavy | 24 |
| 5 | 1 | Scottish Highland | 1: Pony | 50 |
| 5 | 1 | Thoroughbred | 4: Proto-Arab | 20 |
| 5 | 1 | Welsh | 1: Pony | 215 |
| 7 | 2 | Dartmoor | 1: Pony | 46 |
| 7 | 2 | Friesian | 2: Heavy | 9 |
| 7 | 1 | Icelandic | 1: Pony | 10 |
| 7 | 2 | Irish Draught | 2: Heavy | 59 |
| 7 | 5 | Kerry Bog | 1: Pony | 39 |
| 7 | 1 | Lipizzian | 4: Proto-Arab | 42 |
| 7 | 1 | Lusitano | 3: Steppe | 43 |
| 7 | 1 | Maremmano | 4: Proto-Arab | 21 |
| 7 | 1 | Scottish Highland | 1: Pony | 50 |
| 7 | 1 | Viatka | 3: Steppe | 1 |
| 7 | 1 | Welsh | 1: Pony | 215 |
| 8 | 1 | Akhal-Teke | 3: Steppe | 35 |
| 8 | 4 | Andalusian | 3: Steppe | 25 |
| 8 | 2 | Arabian | 4: Proto-Arab | 81 |
| 8 | 1 | Asturcon | 3: Steppe | 13 |
| 8 | 1 | Barb | 3: Steppe | 14 |
| 8 | 2 | Belgian | 2: Heavy | 6 |
| 8 | 2 | Clydesdale | 2: Heavy | 6 |
| 8 | 9 | Connemara | 1: Pony | 58 |
| 8 | 1 | Cukorova | 3: Steppe | 12 |
| 8 | 4 | Dartmoor | 1: Pony | 46 |
| 8 | 6 | Duelmener | 2: Heavy | 10 |
| 8 | 1 | Egyptian | 4: Proto-Arab | 8 |
| 8 | 3 | English Shire | 2: Heavy | 11 |
| 8 | 1 | Friesian | 2: Heavy | 9 |
| 8 | 1 | Fulani | 3: Steppe | 11 |
| 8 | 1 | Garrano | 3: Steppe | 17 |
| 8 | 2 | Guan Mountain | 4: Proto-Arab | 10 |
| 8 | 1 | Iranian | 3: Steppe | 14 |
| 8 | 7 | Irish Draught | 2: Heavy | 59 |
| 8 | 2 | Lusitano | 3: Steppe | 43 |
| 8 | 1 | Merens | 2: Heavy | 12 |
| 8 | 8 | New Forest | 1: Pony | 45 |
| 8 | 1 | Posavina | 2: Heavy | 5 |
| 8 | 1 | Rhineland Heavy | 2: Heavy | 24 |
| 8 | 2 | Shetland | 1: Pony | 64 |
| 8 | 5 | Thoroughbred | 4: Proto-Arab | 20 |
| 8 | 1 | Trakehner | 2: Heavy | 6 |
| 8 | 2 | Vyayskaya | 3: Steppe | 17 |
| 8 | 22 | Welsh | 1: Pony | 215 |
| 8 | 3 | Yakut | 3: Steppe | 21 |

\* indistinguishable from Przelwalskii\_hap14 because missing differentiating site at bp 26

### Phylogenetic relationships based on mitogenomes

The mitogenomes of the 55 newly sequenced Exmoors also were collapsed into eight haplotypes, which corresponded perfectly to the D-loop haplotypes. There were some additional polymorphic sites in the microsatellite repeat region of the control region, but this could have been due to repeat mapping errors and so this region was not considered in the haplotype assignment. Aligning the Exmoor mitogenomes to the sequences described in Achilli et al. (2012) and Kvist and Niskanen (2020) resulted in a total of 94 haplotypes from the 100 published sequences. The phylogenetic tree was highly concordant with the haplogroups suggested by Achilli et al (2012), with high bootstrap support for most internal nodes (Figure 2). The following were not reciprocally monophyletic and so were combined for the purposes of considering relative frequencies: H and I; J and K; O and P; and E, F and G. However, these tended to involve haplogroups that were not well sampled. To allow comparison with Achilli et al. (2012), the tree was rooted with haplogroup R.

**Figure 2.**
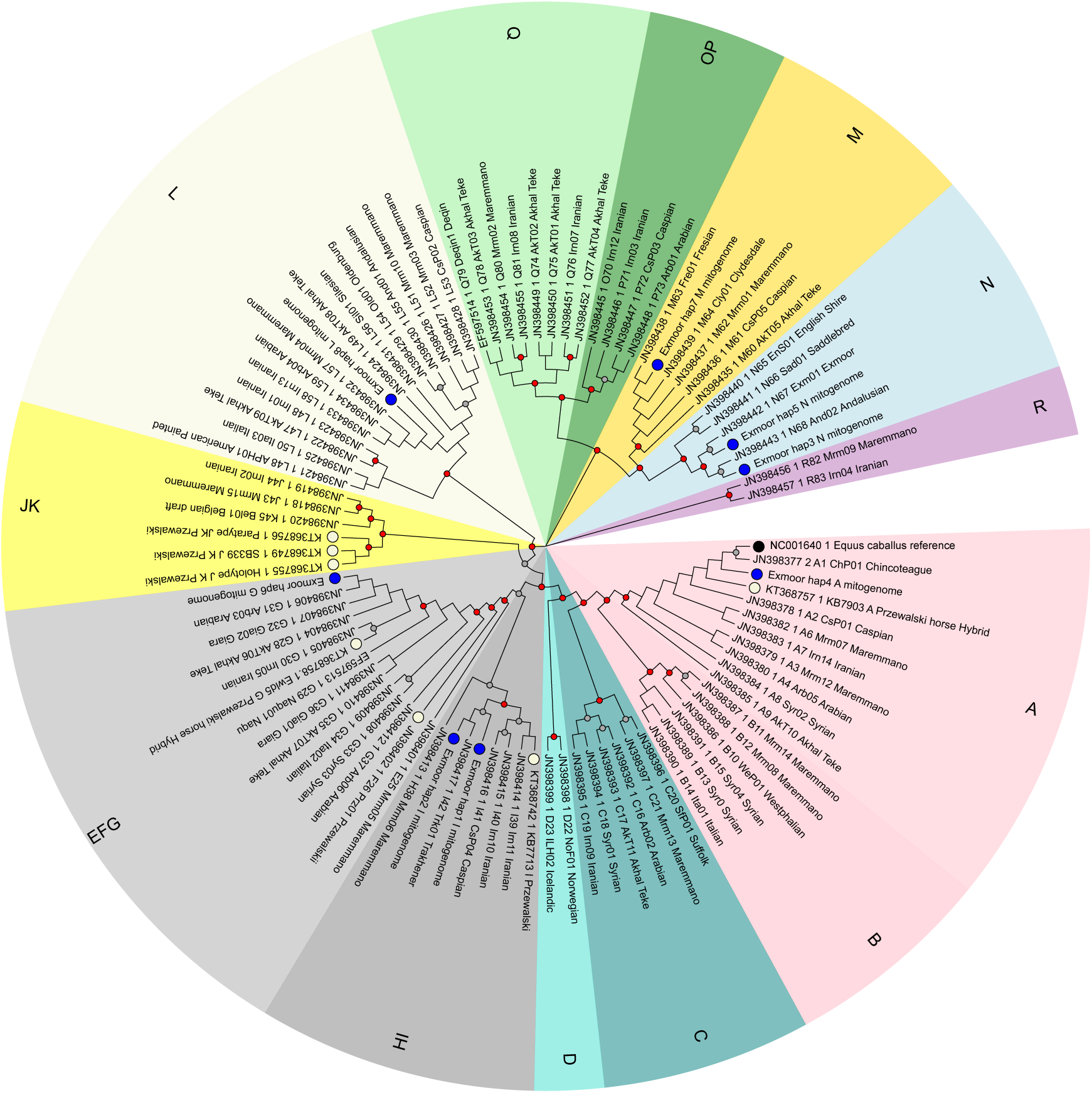
Phylogenetic tree of horse breeds based on complete mitogenome sequences. Letters indicate haplogroups suggested by Achilli et al. (2012), with shading indicating reciprocally monophyletic groups. For example, haplogroups E, F and G would be paraphyletic on their own and so they have been combined based on divergence from the most recent common ancestor that includes all descendents. Circles before the leaf labels indicate groups of interest: blue = Exmoors; beige = Przelwalskii’s horses; black = reference mitogenome sequence of *Equus caballus* (NC001640). As suggested by the haplotype network based only on the D-loop, haplotypes 5 (haplogroup N; identical to the only Exmoor sequence included by Achilli et al.2012) and 7 (haplogroup M) fall into the most basal of the lineages that include Exmoors. The tree is also consistent with Exmoor haplotypes1,2 (haplogroup EFG) and 6 (HI) having unique origins; both haplogroups also include Przewalskii sequences, as does haplogroup A (which includes Exmoor haplotype4 and the *E. caballus* reference genome; this is the most derived clade based on rooting with haplogroup R. Bootstrap support is indicated on the nodes: grey = 70-89%; red = 90-100%; based on the mitogenomes there is strong support for all haplogroup basal nodes, as well as the majority of external nodes.

The Exmoor haplotypes were distributed across haplogroups A (hap4), G (hap 6), I (haps 1 and 2), L (hap 8), M (hap7) and N (haps 3 and 5) (Figures 1 and 2; Supplementary Table 3). The only published Exmoor mitogenome sequence (JN308442; Achilli et al. 2012) was nearly identical to Exmoor_hap5 in our mitogenome sequences and was identical in the D-loop region. No mitogenome sequences from other breeds were identical to the Exmoor mitogenome haplotypes but sequences from Akhal-Teke, Arabian and Iranian breeds (from Achilli et al. 2012) matched Exmoor_hap8 in the D-loop region.

For the 17 Przewalskii horse mitogenome sequences included (two had too many errors to confidently resolve), five mitogenome haplotypes were resolved: three to haplogroup JK (including from paratype and holotype samples; D-loop Przelwalskii hap12); one to haplogroup F (D-loop Przelwalskii_hap11) and another (D-loop Przeslwalskii_hap14) to haplogroup I (Figure 2; Supplementary Table 3). Two sequences from hybrids between Przewalskii and domestic horses (Der Sarkissian et al. 2015) resolved to haplogroups A (with the sequence being very similar to Exmoor_hap4 and the horse reference genome) and G (D-loop Prezwalskii_hap13).

### Phylogenetic relationships based on D-loop sequences

A total of 1060 D-loop sequences (including the 88 newly sequenced Exmoors) were collapsed into 111 haplotypes; an additional 121 “singletons” could not be unambiguously resolved to haplotypes and so these were excluded from the phylogenetic tree (Supplementary Table 3). In general, the D-loop sequences were concordant with the mitogenome haplogroups but with very low bootstrap support (Figure 3). Nevertheless, for most sequences it was possible to unambiguously assign haplogroups. A notable exception was the Przewalskii sequences that had been resolved with high bootstrap support to haplogroup JK in the mitogenome tree (all three mitogenome haplotypes were collapsed to Przewalskii_hap12 for the D-loop only); these appeared in a non-monophyletic (and unsupported) grouping basal to haplogroup L rather than haplogroup JK and were not shared with other breeds. The other Przewalskii sequences were collapsed into Przelwalskii_hap11 (haplogroup F; 15 Przelwalskii sequences, along with three from Mongolian wild horses) and Przelwalskii_hap13 (haplogroup G; two Przewalskii sequences, one Cheju and one Mongolian domestic horse). The Przelwalskii x Domestic horse hybrids were collapsed into Przelwalskii_hap_14 (haplogroup I; including multiple other breeds) and a singleton (haplogroup A) that could not be ambiguously assigned to a haplotype, possibly due to errors in the sequence.

**Figure 3.**
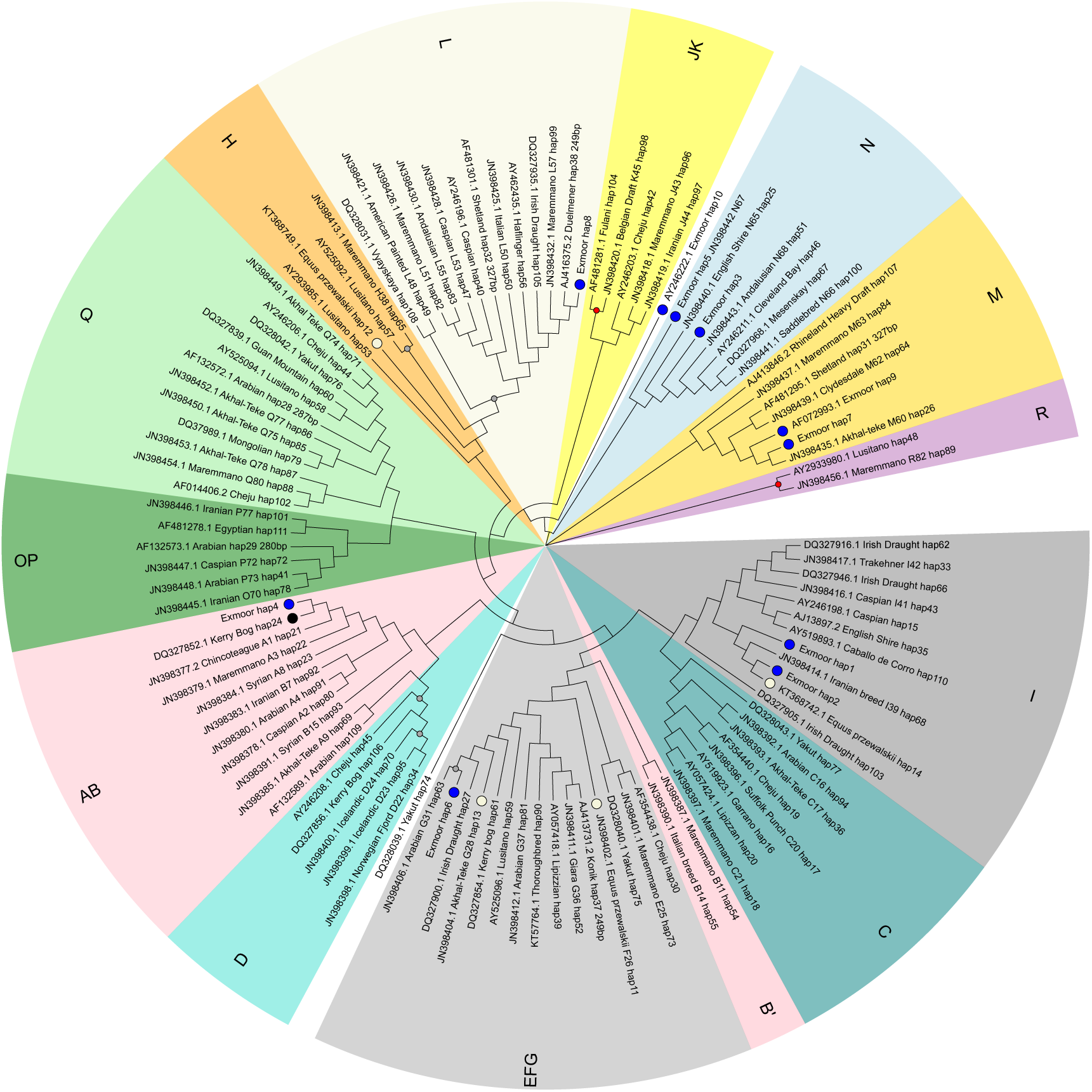
Phylogenetic tree of horse breeds based on D-loop (control region) mtDNA sequences. Letters indicate haplogroups suggested by Achilli et al. (2012), with shading indicating reciprocally monophyletic groups. Circles before the leaf labels indicate groups of interest: blue = Exmoors; beige = Przelwalskii’s horses; black = reference mitogenome sequence of *Equus caballus* (NC001640). Although bootstrap support was generally low (<70%), the D-loop tree resolved most of the mitogenome-based haplogroups, with a few ambiguities: B haplogroup sequences are not monophyletic (some cluster with the A haplogroup, as in the mitogenome tree, whereas those labelled B’ appear as basal to the EFG haplogroup; H haplogroup sequences appear as basal to the L haplogroup but are not resolved). Sequences are named with the following convention: Genbank Accession Number (or Exmoor ID) _Breed_[haplogroup number from the mitogenome tree] _unique haplotype number. Note that two additional Exmoor haplotypes were identified based on published Exmoor sequences but were found in a single sample each: AF072993.1_Exmoor_hap9 was similar to Exmoor_hap7, in haplogroup M; AY264222.1_Exmoor_hap10 was most similar to haplogroup N sequences, but was unresolved.

Haplogroups A and B were also not monophyletic for the D loop, with two haplotypes (haps 54 and 55; labelled as haplogroup B’) appearing as basal to haplogroup EFG (but with bootstrap support less than 50%; Figure 3). Two other sequences were not resolved (hap 74 and Exmoor_hap10); the former included four sequences from the Yakut breed (appearing as basal to the EFG/B’/C and I haplogroups), and the latter was from a single published Exmoor sequence (appearing as basal to the N haplogroup). All other Exmoor haplotypes were clearly resolved into haplogroups.

### Distribution of mitogenome haplogroups across breed groups

Collapsing the 484 Winton et al. (2020) sequences (which started on bp 38 of the alignment) onto the D-loop tree haplotypes resulted in 53 additional haplotypes unique to that set and the rest matched already identified haplotypes, albeit with some important ambiguities (e.g. Exmoor_hap2 vs Przelwalskii_hap14, differed only by a single mutation at bp 26 and so could not be distinguished). However, all haplotypes in the Winton et al. (2020) set could be assigned to mitogenome haplogroups and so they were included in the frequency calculations (Supplementary Table 3). Although there was extensive variation in the number of sequences available for various breeds groups (Supplemental Table 6), the clustering suggested that most have ancestors spanning multiple haplogroups (Figure 4; Supplementary Figures 1-4). However, breeds considered as ponies (including Exmoors) tended to be more distinctive than the other groups, particularly when compared to the southern breeds. Most of the pony breeds included haplotypes in the basal M/N haplogroups that included Exmoor haps 7, 3, 5 and 10 (Figure 4).

**Figure 4.**
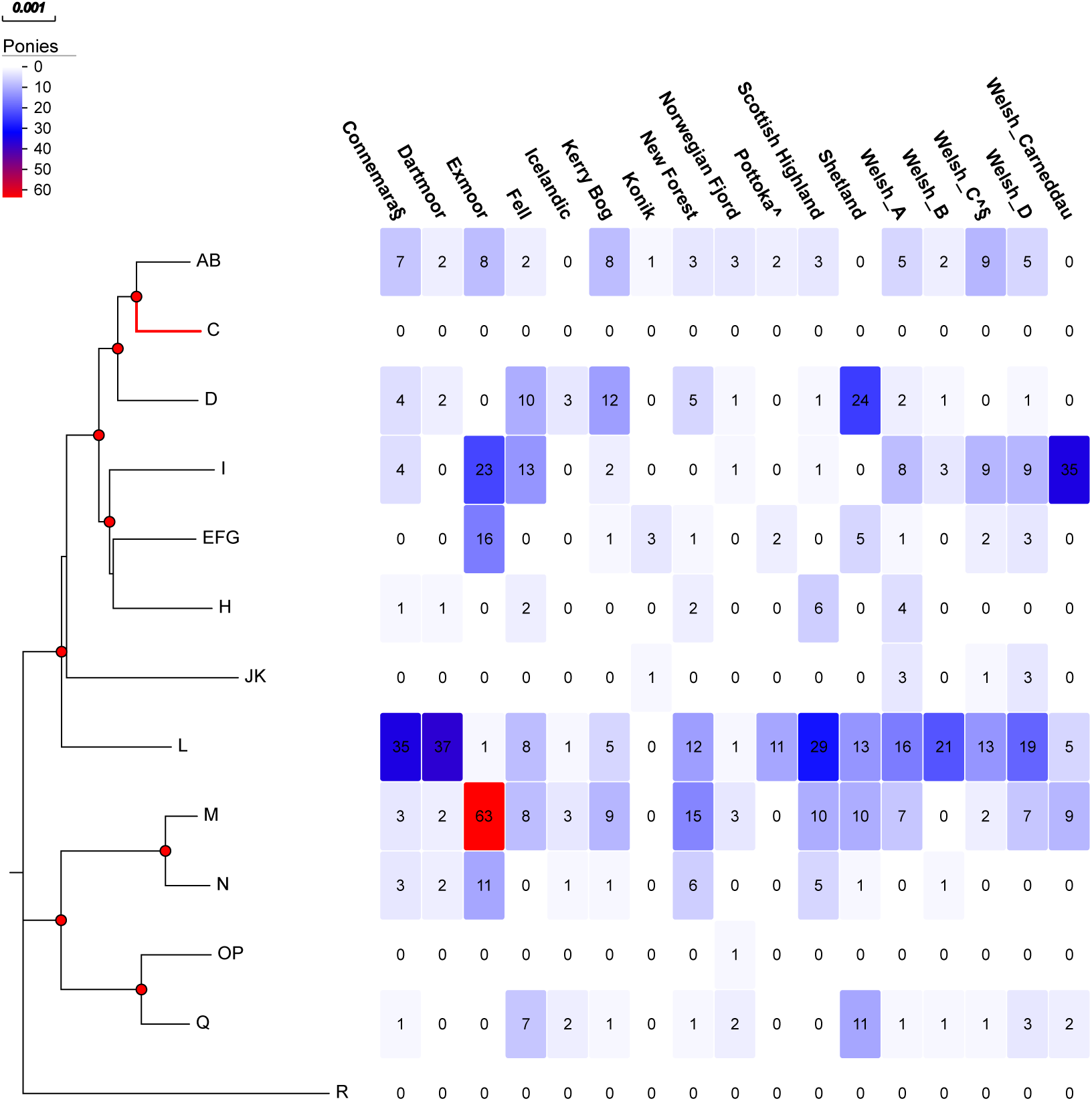
Heatmap showing the relative frequencies of mitogenome haplogroups across various breeds of ponies. The phylogeny on the left shows the haplogroup framework based on a condensed version (one sequence per haplogroup) mitogenome tree (see Figure 2); red circles at nodes indicate bootstrap support >90%). On the heatmap, darker colours indicate higher numbers and red the highest (n = 63). All of the well-sampled pony breeds (except for Pottoka) showed included individuals falling into the Exmoor haplogroups predicted to be ancestral based on both the D-loop haplotype network and the mitogenome tree: haplogroups M (Exmoor haplotypes 7 and 9) and N (Exmoor haplotypes 3, 5 and 10). The I (Exmoor haplotypes1 and 2) and EFG (Exmoor haplotype 6) haplogroups were the next most frequent among Exmoors and shared with some of the other pony breeds. The AB haplogroup (Exmoor haplotype 4) was less common in Exmoors but was represented in most other ponies, except for Icelandic, Shetland and one of the Welsh sections. The L haplogroup was only found in a single Exmoor (Exmoor haplotype 8) but was found in all other well-sampled pony breeds. Haplotypes D, H, JK and Q were found in a subset of the pony breeds, but not in Exmoors. Haplogroups C, H and R were not represented among the ponies and only a single Norwegian sample fell into the OP haplogroup. Although many breeds likely have mixed origins, indicated are those with known influence from: ^ heavy; and § proto-Arab horses.

The exceptions were Konik (n = 5) and Pottoka (n = 15). This is consistent with morphological classifications suggesting that Pottoka has had influence from Type 2 horse breeds, which tend to show a lower frequency of haplogroups M and N than the ponies. Although their highest frequency was found in haplogroups M/N, Exmoors also appear to have had multiple maternal ancestries, with relatively high frequencies of individuals falling into haplogroups AB (Exmoor_hap4) and EFG/I (Exmoor_haps 1,2 / 6), which is consistent with the haplotype network (Figure 1). Connemara (also thought to have influence from Type 4) and Scottish Highland samples tended to show a similar distribution across haplogroups as Exmoors. Haplogroup EFG was shared by some of the other ponies (Kerry Bog, Konik, Pottoka, Shetland and Welsh) but relatively rare in the other breeds. A single Exmoor sequence (hap8) was found in haplogroup L, which is at high frequency in breeds distributed across all morphological breed groups. Ponies were not represented in haplogroups C, H or R and OP was found in only a single individual from one breed (Norwegian Fjord); JK was only found in Konik and Welsh breeds. Haplogroups D and Q were found in Connemara, Fell, Icelandic, Kerry Bog, New Forest, Norwegian Fjord, Shetland and Welsh sequences but neither was found in Dartmoor, Exmoors, Konik, or Pottoka.

The northern horses (Type 2: Cold-blooded or Heavy) were not as well sampled as the ponies but their distribution across haplogroups was not dissimilar: haplogroups L and AB were most represented, followed by I/EFG and M/N. However, all haplogroups were represented by at least a single sequence from a single cold-blooded horse breed (Supplementary Figure 2).

The other recognised ancient lineage of horses (Przelwalskii) has at least part of their ancestral origins as Steppe horses, in the southern breed group (Supplementary Figure 3). Notably, none of the Przelwalskii sequences were resolved to haplogroup L (except for hybrids with domestic horses) but there appeared to be two clusters of ancestral types: I/EFG (similar to Exmoors) and JK (which is relatively rare in other breeds and breed groups). Among the other Steppe horses, haplogroup L was again most frequent, followed by haplogroup AB and then I/EFG. Haplogroup Q was more common in this breed group (but not in Przelwalskii’s). All haplogroups were represented among more than one Steppe breed and some breeds (e.g. Akhal-Teke) were widely distributed across haplogroups.

For the other southern horses (Proto-Arab), Arabian horses were well sampled (n= 82) and showed a high frequency of sequences in haplogroups I/EFG, AB, L, OP, and Q with only four in haplogroup M/N and one in haplogroup R. Haplogroups D, H, and JK were not represented in the Arabian haplotypes. Not many thoroughbreds were sampled in the focal studies (n= 20, including the *Equus caballus* reference genome) but they were at highest frequency in haplogroup L, with scattered sequences in other haplogroups but not AB, C, JK, OP, Q or R.

### Persistence of haplogroups over time

All of the “modern” mitogenome haplogroups were represented among the sequences from the ancient DNA samples compiled by Cielsnak et al. (2010) but there were also four clusters that were only found in the ancient samples (haplogroups S-V; Supplemental Table 6). Haplogroup V was only found in a single late Pleistocene/Mesolithic sample from the Iberian Peninsula and so was not considered in the frequency analysis of changes over time (Figure 5). Only the Iberian Peninsula was sampled across all time periods (n = 33; Supplementary Table 6), starting in the Mesolithic (5380-4900 BC), so only the major time periods (rather than geographic locations) are considered in relation to the backbone haplogroup phylogeny (Figure 5). Haplogroup S was only found in Pleistocene samples from Alaska and Siberia, whereas haplogroup T was found in all time periods until the Bronze Age, across most geographic regions. Haplogroup U was rarer but found in Neolithic and Bronze Age Iberian samples and an Iron Age sample from China.

**Figure 5.**
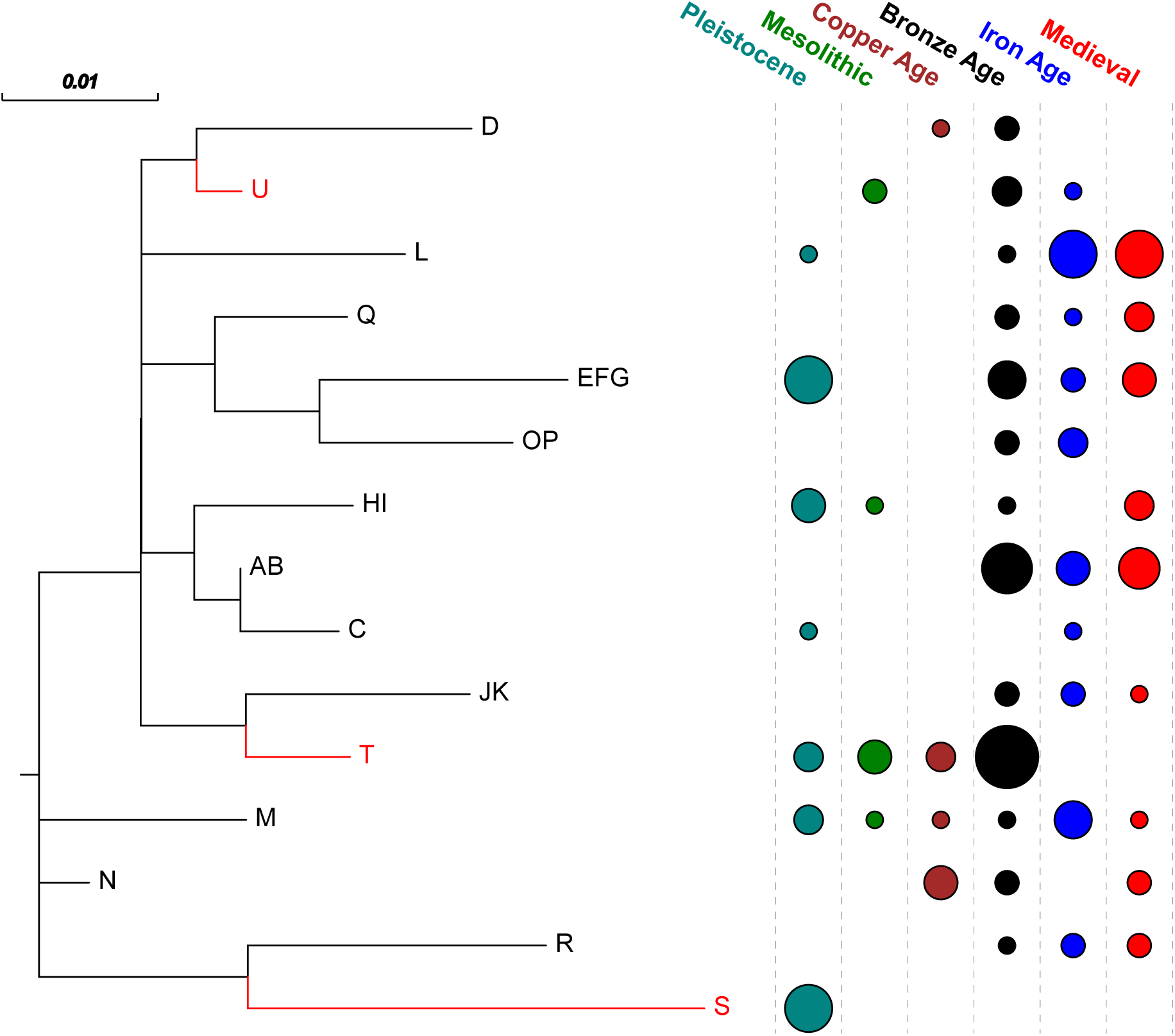
Backbone D-loop phylogenetic tree including ancient DNA sequences, using the samples used to create the mitogenome framework tree, along with one sequence from each of the haplotype groups that were only found in ancient sequences (which were assigned arbitrary letters S-U, branches and labels in red) and so may be extinct. The tree was largely unresolved (with no bootstrap proportions >70%) with respect to relationships among haplogroups but provides a framework for assessing persistence of lineages over time. Only two samples (sourced from Europe, Asia Minor, Armenia) represented the Enolithic era (not shown here) but they clustered to haplogroups HI and M.

Although the later time periods tended to be more widely sampled (Bronze Age: n=45; Iron Age: n =67; Medieval: n = 30) than the earlier (Pleistocene: n = 26; Mesolithic: n = 8; Copper Age: n = 9), there are some interesting patterns in relative persistence of haplogroups (Figure 5). Haplogroup M (hypothesised to be the ancestral Exmoor pony haplogroup) persisted across all time periods and across regions sampled. Haplogroup L was also persistent but not distributed as widely geographically (Supplementary Table 6) and most frequent among Iron Age and Medieval samples. With the caveat of limited sampling in the earlier time periods, haplogroups AB (frequent among all pony and horse breed groups), JK (only found in Konik and Welsh modern pony breeds but more common among southern horse breed groups), OP (not found in modern ponies or heavy horses except for single Norwegian Fjord and Rhineland heavy horse samples but frequent in southern breed groups), Q (frequent in some modern ponies and southern horse breed groups but not found in Exmoors), and R (found in southern horse breed groups but not ponies or northern horses) may have first appeared in the Bronze Age. However, haplogroup JK shares a recent ancestor with the “extinct” T haplogroup and haplogroup R shares an ancestor haplogroup S, suggesting possible divergence of the ancestral lineages over time. Haplogroup D, which was frequent in modern ponies (but not found in Exmoors), was rare among the ancient samples but found as early as the Copper Age. Haplogroup C, which was not found in modern ponies but represented among all three horse breed groups, was rare in the ancient samples but appeared in European samples from the late Pleistocene and was the only haplogroup (except for S) not identified among the Bronze Age samples.

## Discussion

Overall, our study emphasises the value of long-term studbook records to inform conservation management (Moreno et al. 2024), particularly when combined with genetic perspectives based on sequence data to determine relationships among historical lineages. In contrast to other studies suggesting that the D-loop does not capture enough variation to confidently resolve haplotypes (Achilli et al. 2012), our sequencing of the mitochondrial D-loop, representing most of the lineages described in the Exmoor pony stud book (n = 88 samples, from 24 maternal founder lines), captured all variation determined by full mitogenome sequencing. A recent meta-analysis (Agbani et al. 2025) suggested that existing frameworks for classifying horse haplotypes do not sufficiently capture evolutionary relationships but we found that the mitogenome framework of Achilli et al. (2012), with a few exceptions, seems robust and allows a useful perspective to describe maternal diversification patterns, including sequences from ancient DNA samples. To resolve relationships among lineages, it would be useful to obtain additional mitogenome sequences from those that were ambiguous just based on the D-loop, as recommended by Agbani et al. (2025). However, if the purpose is to identify haplogroups and predict shared ancestry, the D-loop appears to capture much of the variation. Our combined analysis of D-loop, mitogenome and ancient DNA sequences thus provides a platform to assess hypotheses about maternal origins of modern pony breeds, including Exmoors.

### D-Loop Exmoor haplotype distribution across maternal lineage and other breeds

Although the D-loop sequences revealed some errors in the information provided with the samples, clustering of founder lines by mtDNA haplotype mostly aligned as expected according to the location and management history of the foundation matrilines, indicating that most recording of maternal pedigrees in the Exmoor Pony Stud book has been accurate, even for free-living herds, as has been found in similar studies of Soay sheep on St. Kilda, where observation records are comprehensive (Bérénos et al. 2014; Hunter et al. 2019). For breeds with strict registration procedures such as Exmoor ponies, our results thus reveal the value of mtDNA sequencing to verify maternal lineages, as has been suggested for other breeds such as thoroughbreds (Hill et al. 2002). However, our results also emphasise risks of relying on published databases: although two additional D-loop haplotypes were resolved in Exmoors from published sequences, these were based on single sequences from unpublished Genbank entries and so could represent sequencing or studbook assignment errors; one (hap_9; AF072993: Lister et al., unpublished) was only a single bp different from Exmoor_hap7 but the other (hap_10; AY246222: Flannery and Cothran, unpublished) was quite distinctive and unresolved in the D-loop phylogeny. In addition, our Exmoor_hap8 was only found in a single Exmoor sample but was frequent in other breeds; since this was the only sample included from its ancestral maternal lineage (76003) further investigation would be required to verify the provenance of this dam line. While it would also be valuable to use Y chromosome markers to verify paternal ancestry, the Y chromosome of horses has very low levels of polymorphism (Lindgren et al. 2004; Lau et al. 2009; Cieslak et al. 2010), reducing this potential. We did attempt to map our whole genome sequence data to the Y chromosome but there was not enough variation to distinguish between paternal lineages (Davy 2024).

Our study revealed three additional haplotypes not identified in other studies that included Exmoors, suggesting a broader range of maternal origins of Exmoors compared to the previously most comprehensive study on British breeds (McGahern et al. 2006). The 20 Exmoor sequences they included were resolved into only two of seven British clades. The very long branches separating the additional haplotypes that we found, even based on 350 bp of D-loop sequence, could suggest a more complicated maternal history.

More diverse origins than studbook predictions also has been found for other rare British horse breeds, such as Cleveland Bay; one of only five domestic horse breeds, (including Exmoors), classified as Critical on the Rare Breeds Survival Trust UK Watchlist (Rare Breeds Survival Trust 2025b). This breed had been thought to be founded by a single extinct ancestral type but was found to have had at least four maternal ancestry lines based on D-loop variation (Dell et al. 2020). Although they didn’t include Exmoors, Winton et al. (2020) sampled British and Irish pony breeds more widely and compared mitochondrial with nuclear diversity based on microsatellites; they also concluded complex histories but maintenance of rare maternal lines in some of the breeds, along with different histories of maternal and paternal influences, with the latter including a wider range of non-pony breeds. They found that Welsh ponies had more haplotypes than other ponies (n=42) but sampling was also more extensive than for the other breeds. Nevertheless, identification of distinct ancestral lines could have important implications for conservation of particularly rare lineages within these breeds.

Although only informative about maternal origins, our results suggest that there could have been differential mixing between different Exmoor lineages with other breeds. Of the 87 included in our study, 37 other breeds shared Exmoor haplotypes (Supplementary Table 7) but the relative frequency differed across lineages. The haplogroup I lineages (Exmoor_haps1 and 2; found in single maternal lines from foundation herds 12 and 76, respectively) showed very limited sharing with other breeds, although they are highly similar to haplotypes found in Przelwalskii horses (Exmoor_hap2, only differing by a single bp from hap14 in the D-loop). This could suggest that these herds represent historic lineages that have been kept isolated from other breeds. Interestingly, individuals with Exmoor_hap1 were all sampled from Exmoor, whereas those from Exmoor_hap2 were more broadly distributed (Exmoor, Lancashire, Brecon Beacon, Fort Augustus, Lancashire, Scorraig), consistent with isolation from other breeds in the former but some mixing in the latter. Exmoor_hap6 (haplogroup G) was also only found in a single maternal line from foundation herd 12 and was not found in other breeds, but most of the samples were from individuals that had died, except for two individuals that had been moved to Scotland. This suggests that this rare and unique lineage is at risk of extinction; however, the surviving animals from this maternal line have recently been recruited into a rewilding project (Bamff Wildlands) originally focused on beavers (Law et al. 2017) but now being extended to include terrestrial macroherbivores. Given the retention of ancestral traits for environmental resilience exhibited by Exmoors (Gates 1979; Baker 2008; Hovens and Rijkers 2013), using such projects could provide a platform for preserving the breed in the long term.

The A/B haplogroup lineage (Exmoor_hap4) from a single maternal line in foundation herd 23 was only shared with other ponies but was nearly identical in the D-loop to the horse reference sequence; whether this reflects hybridisation of Exmoors with thoroughbred mothers, or pony ancestry of thoroughbreds would require further investigation. Exmoor_hap3 was only found in a single individual but it was also the only representative from foundation herd 44; although also rare, the fact that it was shared with heavy horse breeds also could suggest mixing between different breed groups. Whole genome sequencing would be useful to more clearly distinguish ancestral origins from subsequent admixture.

One of the Exmoor haplotypes (Exmoor_hap 7; haplogroup M) was found in over half of the dam lines sequenced but this could be due to the history of the founder lines. The original anchor herd on Exmoor (900) only included this haplotype and herd 01 included haplotypes 5 (haplogroup N) and 7, consistent with conclusions based on the haplotype network that these are the more ancestral lineages. Shared ancestry of herds 900 and 01 is also in line with evidence that these herds were closely connected during the 1950s (Baker 2008). These haplotypes also were shared across breed groups, but mostly only in one or two samples from the other breeds (except for Kerry Bog, which had Exmoor_hap7 in 13% of individuals sampled). The relative rarity in other breeds and high frequency in Exmoors could suggest that these Exmoor lineages have contributed to a wide range of other breeds, as suggested by (Baker 2008). These lineages also might have contributed to foundation of later Exmoor herds, as Exmoor_hap5 was also found in a maternal line from herd 23 and Exmoor_hap7 in one from herd 48. Perspectives from whole genome sequencing could be used to test these hypotheses more thoroughly (Davy 2024).

The mtDNA results also might point out errors in the registration of individuals for the studbook records. For example, in contrast to the other haplotypes, Exmoor_hap8 (haplogroup L) was only found in a single Exmoor individual (from herd 76) but was frequent (n=98 samples) across all breed groups, including other ponies (n=45). Since this was from a maternal lineage accepted into the Stud Book based on phenotype evaluation rather than recorded pedigree and the parents of the mare (born in 1960, just before the studbook closed) were not recorded, it is possible that it does not represent a “true” Exmoor individual but reflects maternal contributions from other breeds.

### Phylogenetic relationships based on mitogenomes

Our mitogenome analysis further supported a complex evolutionary history of Exmoor maternal lineages. The framework proposed by Achilli et al. (2012) was generally well resolved, except for haplogroups that had not been well sampled (haplogroups E, F, H, K, O) and did not appear as reciprocally monophyletic and so were collapsed into a smaller number of haplogroups (EFG, HI, JK, OP). Achilli et al. (2012) had only sampled a single Exmoor, which was identical to our Exmoor_hap5 (haplogroup N) but our mitogenome sequences suggest four potentially distinct origins: Exmoor_hap4 (haplogroup A, including the Equus caballus reference genome and thoroughbreds); Exmoor_haps1/2 and 6 (haplogroup EFG and HI, both including Przelwalskii’s horses); Exmoor_haps3/5 and 7 (haplogroups N and M); and Exmoor_hap8 (haplogroup L, including a wide range of breeds). Based on relative sequence divergence, we rooted the mitogenome tree using haplogroup R, which would suggest that the N/M Exmoor haplotypes are the most ancestral and haplogroup A the most derived, which is consistent with our haplotype network and with predictions based on population and herd histories plus studbook pedigrees. Sampling only Exmoors would have made it more challenging to predict ancestries, without the broader context based on other breed groups.

The origin of Przelwalskii’s horses (thought to be extinct in the wild since the 1960s) has been controversial in relation to admixture with domesticated horses (Ishida et al. 1995; Lau et al. 2009; Der Sarkissian et al. 2015) but our results suggest that they have had different ancestry from the foundation Exmoor herds. Achilli et al. (2012) had only included two Przelwalskii’s horses (both resolved into mitogenome haplogroup F; D-loop Przelwalskii_hap11, shared only with Mongolian domestic horses) but inclusion of additional mitogenomes (Der Sarkissian et al. 2015) suggests the possible influence of three maternal lineages: haplogroups JK (paratype and holotype: D-loop Przelswalskii_hap12), G and I. Since only Przelwalskii_hap13 (haplgroup G) was shared with non-Steppe horse breeds, this could reflect admixture of this maternal lineage with domestic horses rather than shared ancestry. Interestingly, the Exmoor haplotypes in the haplogroups including Przelwalskii’s were only found in single herds: Exmoor_hap1 and 6 in herd 12 and Exmoor_hap2 in herd 76. It is thus possible either that some of the Exmoor herds had maternal influence from Przelwalskii’s or that they shared a common ancestor with other domesticated breeds, which would be consistent with the whole genome analyses of Der Sarkissian et al. (2015) suggesting extensive admixture of Przelwalskii’s with domesticated horses since their diversification ∼45,000 years ago.

### Phylogenetic relationships based on D-loop sequences

Although there was much lower resolution based on bootstrap support when considering only D-loop sequences, as noted in other studies (Vilà et al. 2001; Jansen et al. 2002; Royo et al. 2005), there was still remarkable concordance with the mitogenome haplogroup framework. While some of the mitogenomes were collapsed into single D-loop haplotypes it was still possible to map even truncated sequences (for which individual haplotypes could not be assigned with confidence) to mitogenome haplogroups. However, there were some additional ambiguities in relationships among haplogroups suggesting that resolution of the deeper nodes may not be reliable just based on the D-loop. Notably, D-loop sequences that would have been assigned to haplogroup H based on their mitogenome sequences (e.g. Przelwalskii horse holotype/paratype) did not cluster with haplogroup I and instead appeared as basal to haplogroup L (albeit with no bootstrap support). Haplogroups A and D were also paraphyletic (i.e. not all descendants of the predicted shared ancestor were included in their groupings) and B was both paraphyletic with A and polyphyletic (with two sequences that would have been assigned as haplogroup B based on their mitogenomes appearing as basal to EFG rather than clustering with A). The mitogenome phylogeny suggests that haplogroups A, B, C and D share a common ancestor and so could be considered a single lineage but the D-loop phylogeny suggested that haplogroup C did not share an immediate ancestor with AB and D and were more closely related to haplogroup I. This could affect interpretation of origins of breeds that fell across these haplogroups based on the D-loop tree but did not have mitogenomes included. For example, for Kerry Bog, which McGahern et al. (2006) resolved into six clades across their D-loop phylogeny (based on the same 39 sequences that we included), we resolved 16 haplotypes that spanned eight haplogroups, including A, B, D and I. Thus, variation in D-loop sequences on its own may not have enough power to resolve deeper relationships among haplogroups but can still be useful to identify more recent relationships and diversity of maternal origins among breeds.

Cieslak et al. (2010) found that phylogenetic resolution was not substantially increased by comparing the entire mitochondrial control region and suggested that sequencing of a larger proportion of the mitogenome might be necessary; however, lack of resolution might also be due to the large number of maternal lineages found within breeds. A large number of female founders is consistent with the sex-biased dispersal and breeding of domestic horses throughout their history. It is usual for many females to mate with one male, leading to a much larger female effective population size, and therefore higher levels of diversity in the maternally inherited mitochondrial genome compared to what might be found in the nuclear genome (Vilà et al. 2001). This means that analysing mtDNA alone can lead to an overestimation of the genetic diversity within populations, even if based on the entire mitogenome.

### Distribution of mitogenome haplogroups across breed groups

Although the maternal ancestry of mtDNA means that it is not informative about cross-breeding among ancestral breed groups, our mapping of the morphologically designated breed groups proposed by Schafer (1981) is consistent with the complex histories of horses. Although there is a large bias in the representation of various breeds based on published sequences, nearly all well-sampled breeds (including Exmoors) included D-loop haplotypes that fell across multiple mitogenome haplogroups, which is expected given the known mixing among modern breeds, including ponies (Winton et al. 2020). However, there did appear to be some distinction in haplogroup frequency patterns between northern (ponies and heavy horses) and southern breed groups, again consistent with the morphological predictions (Schafer 1981). Discrepancies between maternal and nuclear phylogenies could help to identify both past and recent admixture. For example, a nuclear DNA study by Peterson et al. (2013) based on 10,066 SNPs from the Illumina 50K SNP beadchip, found that individual breeds formed distinct monophyletic clades in a phylogeny of 814 sequences from 38 domestic horse breeds, and that the majority of breeds clustered in accordance with the ancestral groupings described by Schaffer (1981), including Exmoor ponies, which clustered with Type 1 pony breeds. This emphasises that whole genome sequencing could be used to make more robust inferences about admixture and origins of extant populations (Davy 2024).

### Persistence of haplogroups over time

Distribution of haplogroups in the ancient samples generally supported predictions based on the modern haplogroup frequencies in relation to ancestry of ponies and distinctive origins of northern and southern breed groups. Although the R haplogroup inferred to be basal in the mitogenome phylogeny was not identified in the Pleistocene samples, the apparently extinct S haplogroup was closely related to this and appeared as basal in the backbone phylogeny that included the ancient DNA samples. It is intriguing that haplogroups frequent in the southern breed groups (Steppe and Proto-Arab) but rare in the northern breed groups (ponies and cold-blood horses) were not represented in pre-Bronze Era ancient DNA sequences. This is consistent with the view of Schafer (1980) that northern and southern lineages had different ancestries.

The persistence of the M haplogroup across time periods (containing haplotypes found in the anchor Exmoor herds) is also consistent with the view that Exmoors represent an ancient lineage, little changed from the primitive ‘Universal Pony’ (Speed and Speed 1977; Schafer 1981; Baker 2008). An exciting area of further study would be a comparison, using molecular methods, of the modern Exmoor population with further carbon-dated ancient samples (and samples of modern breeds currently lacking published data) originating from the UK and Western Europe, to investigate the validity of these assertions.

## Conclusions

Overall, our results are consistent with the view that the history of horses, before the advent of selective breeding by people, is complex. We suggest that existing mtDNA sequencing studies provide a wealth of data that could be interpreted more clearly in the context of the haplogroup framework proposed by Achilli et al. (2012). Although comparison of nuclear genomes would be necessary to resolve predictions about admixture, we focused on mtDNA here because of the amount of data already available. Even though we included D-loop sequences of varying lengths, mapping sequence variants onto the mitogenome structure allowed testing of hypotheses about the diversity and origins of putatively ancient domestic breeds. Additional mitogenome sequences would be required to resolve some of the deeper relationships among haplogroups but many lineages are already well represented. The approach we used to assess diversity in relation to what is known about pedigrees and management histories of the Exmoor pony can be applied to other rare or ancient breeds (that have been sampled comprehensively enough).

Our focus on the Exmoor pony was because of its exceptional retention of survival traits, the increasing interest in its performance as an eco-engineer, and the extensive pedigree records available to inform selecting a representative sample. In addition, the still debated hypothesis of its ancient origins and primitive status needed to be resolved as a priority, given the considerable implications for the conservation of the Exmoor pony population. Our broad sampling across maternal lineages traced the most basal lineages of Exmoors to the Pleistocene, making a substantial contribution in support of its hypothesised antiquity. We identified new haplotypes that had not previously been reported and identified basal haplotypes that may need to be prioritised for conservation. Another important next step would be to assess whether the different haplogroup origins of Exmoors also represent distinctive morphologies, including preservation of traits for resilience that make them so effective for rewilding natural environments.

## Acknowledgements

We thank Anubhab Khan for training in bioinformatics for the mitogenome mapping. We thank the Exmoor Pony Society for funding DD and providing access to the EPS studbook and the Rare Breed Survival Trust for access to additional records.

## Author Contributions

DD designed the study, collected the samples, generated the pedigrees and contributed to writing and interpretation. AH, AB and FA-G contributed to data analysis and writing. EK performed all of the lab work to generate the molecular data. SB provided access to historical archives and contributed to writing. PD contributed to writing. BKM took the lead on data analysis and writing and supervised the work of students contributing to the paper.

## Data Archiving

The Exmoor pony mitogenome sequences assembled based on short-read sequences for this study have been deposited to Genbank (accessions numbers pending). Details about the relative frequency of haplotypes is provided in the supplementary information. Alignment files will be available on the University of Glasgow Enlighten database (access number pending).

