## Supplementary Table captions and Figures for "Origins and diversity of Exmoor ponies: A mitogenome framework for assessing maternal lineage diversification in endangered pony breeds"

### Supplementary Tables and Figures

**Supplemental Table 1. Details of Exmoor pony individuals sequenced in the study**, indicating their data from the Exmoor pony society studbook records (individual code, whether they were living or dead, their gender, date of birth (DoB), sire and dam found lines), D-loop haplotype, mitogenome haplogroup assigned, and whether they had a mitogenome sequenced. Locality data are not included, to protect the privacy of the owners who donated samples.

**Supplemental Table 2. Summary statistics of whole genome sequencing of Exmoor pony samples (N = 55)**, showing the individual code, the sequence batch date, the total number of raw reads, average whole genome-wide coverage, mitochondrial genome coverage, and the number of reads mapped to the mitochondrial genome.

**Supplementary Table 3. Details of D-loop sequences included in the study,** indicating: accession numbers for published sequences or individual codes for newly sequenced Exmoor samples; the breed assigned by the submitting authors; the length of the aligned region; the D-loop haplotype; the mitogenome haplogroup; whether a mitogenome was included in the mitogenome phylogeny; the source; whether the sequence was included in the synthetic studies by McGahern et al. 2006 and Kvist and Niskanen 2020; and the morphological breed group assigned. Singletons are sequences only found in a single individual but for which a haplogroup could be assigned based on clustering; these were not included in the phylogenetic analyses, except for mitogenome singletons from Achilli et al. (2012) because they only included unique haplotypes in their study.

**Supplementary Table 4. Ancient DNA sequences compiled by Cielsnak et al. 2010,** indicating the geographic region, timings, geographic location and dates assigned in that study, along with the original source and the haplogroups that we assigned based on clustering analyses (excluding those . We excluded sequences that could not be confidently resolved to haplogroups.

**Supplementary Table 5. Variation among Exmoor D-loop haplotypes,** showing the position in the 350 bp alignment of polymorphic sites, frequency in newly sequenced (new = 88) and published (old = 32) Exmoor samples, along with the number of sequences and list of other breeds sharing individual haplotypes (in parentheses). Note that the Welsh samples (from Winton et al. 2020) listed next to haplotype 2 could be Przelwalskii_hap14 because they are missing the region including a distinguishing SNP at the start of the sequence. Yellow highlighting indicates mutations restricted to one haplotype; purple indicates shared polymorphisms.

**Supplementary Table 6. Haplogroup frequency of ancient DNA samples compiled by Cieslak et al. 2010**, indicating the time period, the estimated date ranges (where specified), and assignment to the mitogenome haplogroups. Sequences in haplogroups S, T, U and V did not cluster with the mitogenome haplogroups from extant sequences and so were assigned to new haplogroups. Haplogroup V was only found in a single sample (from a late Pleistocene/Mesolithic sample from the Iberian Peninsula) and so was excluded from the frequency analysis in Figure 5.

**Supplementary Table 7. Number of D-loop haplotypes resolved relative to the total number of** sequences for 88 extant breeds of horses, indicating the ancestral breed group assigned and whether the breed shared Exmoor or Przelwalskii haplotypes.

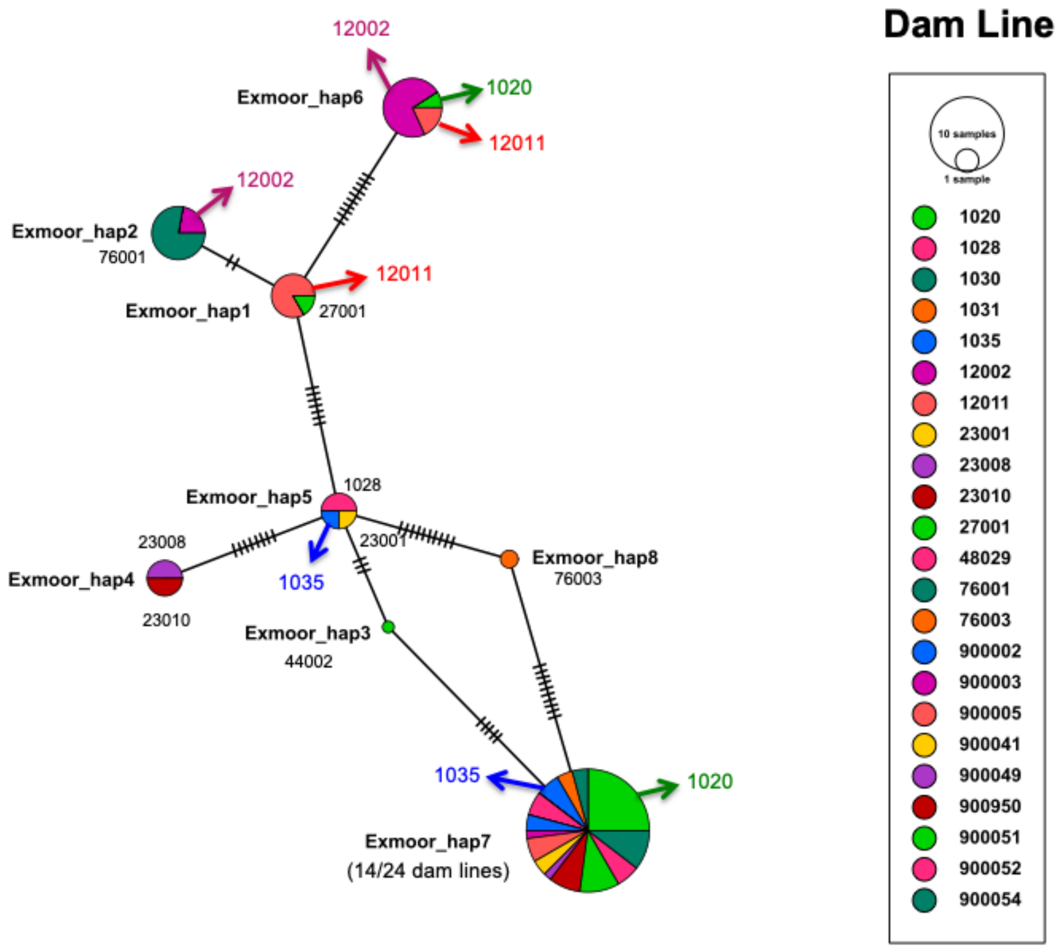


**Supplementary Figure 1. Minimum spanning D-loop haplotype network of newly sequenced Exmoor pony sequences, grouped by maternal founder lines (Dam lines), prior to error detection.** Size of the circles is proportional to the sample size; colours indicate maternal lineages. Hatch marks indicate the number of mutations separating each haplotype. Arrows indicate maternal lineages that included more than one mtDNA haplotype, suggesting discrepancies in the studbook records or metadata associated with the sample. In all cases, matrilines with multiple lineages were due to naming errors on the samples, rather than errors in the studbook record.

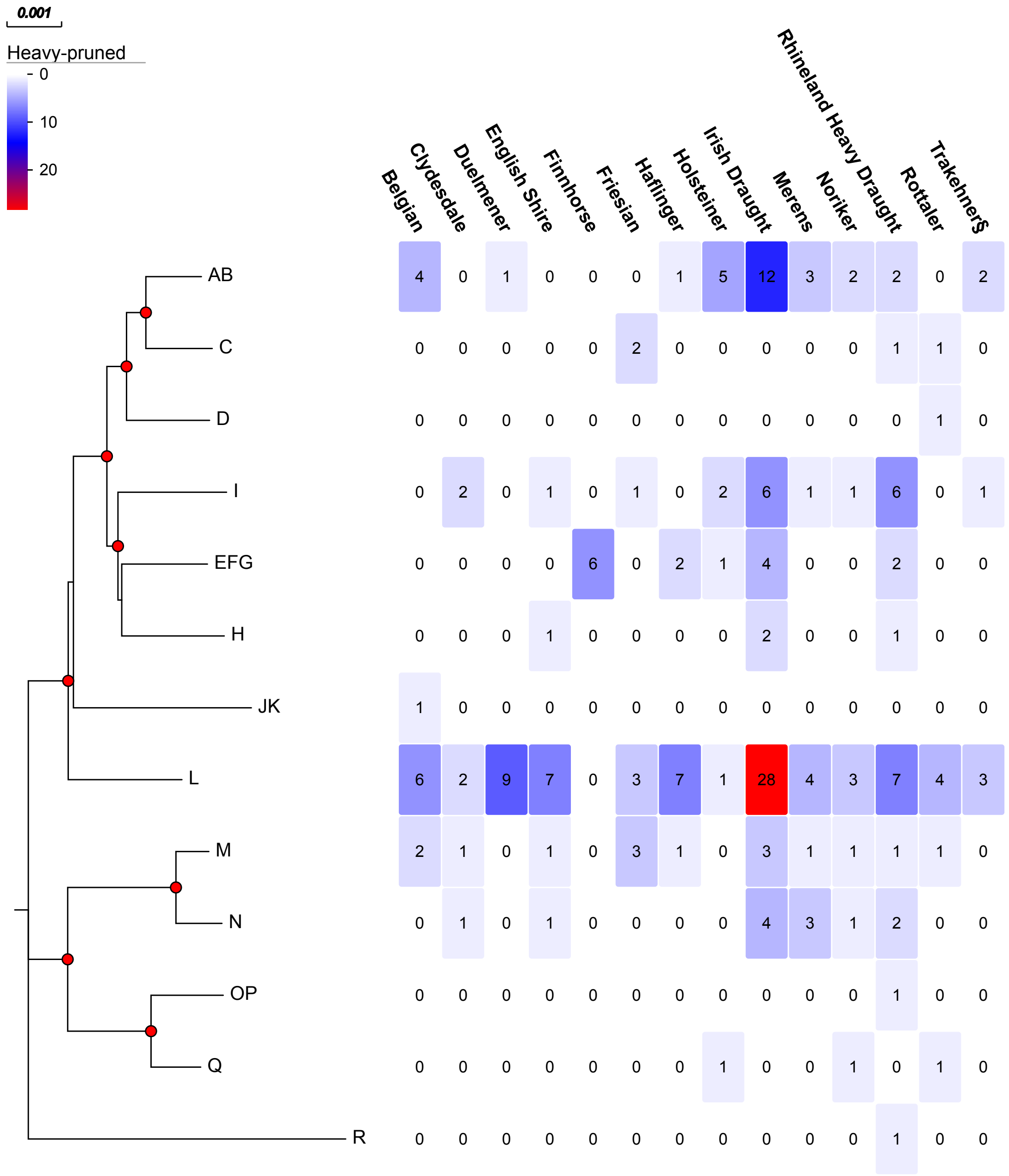


**Supplementary Figure 2. Heatmap showing the relative frequencies of mitogenome haplogroups across horse breeds classified as Type2: Cold-Blood (Heavy), visualised using Evolview v2 (He et al. 2016).** The phylogeny on the left shows the haplogroup framework based on a condensed version (one sequence per haplogroup) mitogenome tree (Figure 1); red circles at nodes indicate bootstrap support >90%. Each column on the right is a breed classified as Cold Blood or Heavy, in the Northern Group horses (Schafer 1980), showing the number of sequences from each haplogroup. Only breeds with at least 5 sequences are included (see Supplementary Table 2 for full classifications). Note that most breeds (except Finnhorse, which only had 6 sequences available) had representatives in the L haplogroup but the AB and I haplogroups frequent in the pony breeds (see Figure 4) were also well-represented. Although the M and N haplogroups were represented in some of the Heavy breeds, they tended to be less frequent than in the pony breeds. The EFG haplogroup was also less common among the Heavy than the pony breeds.   The C, H haplogroups not found in the pony breeds were represented among some of the heavy breeds.  Although rare, there were also sequences resolved to the D, JK, OP and R haplogroups.  Although many breeds likely have mixed origins, indicated are those with suspected influence from: ^ heavy; * steppe; and § proto-Arab horses.

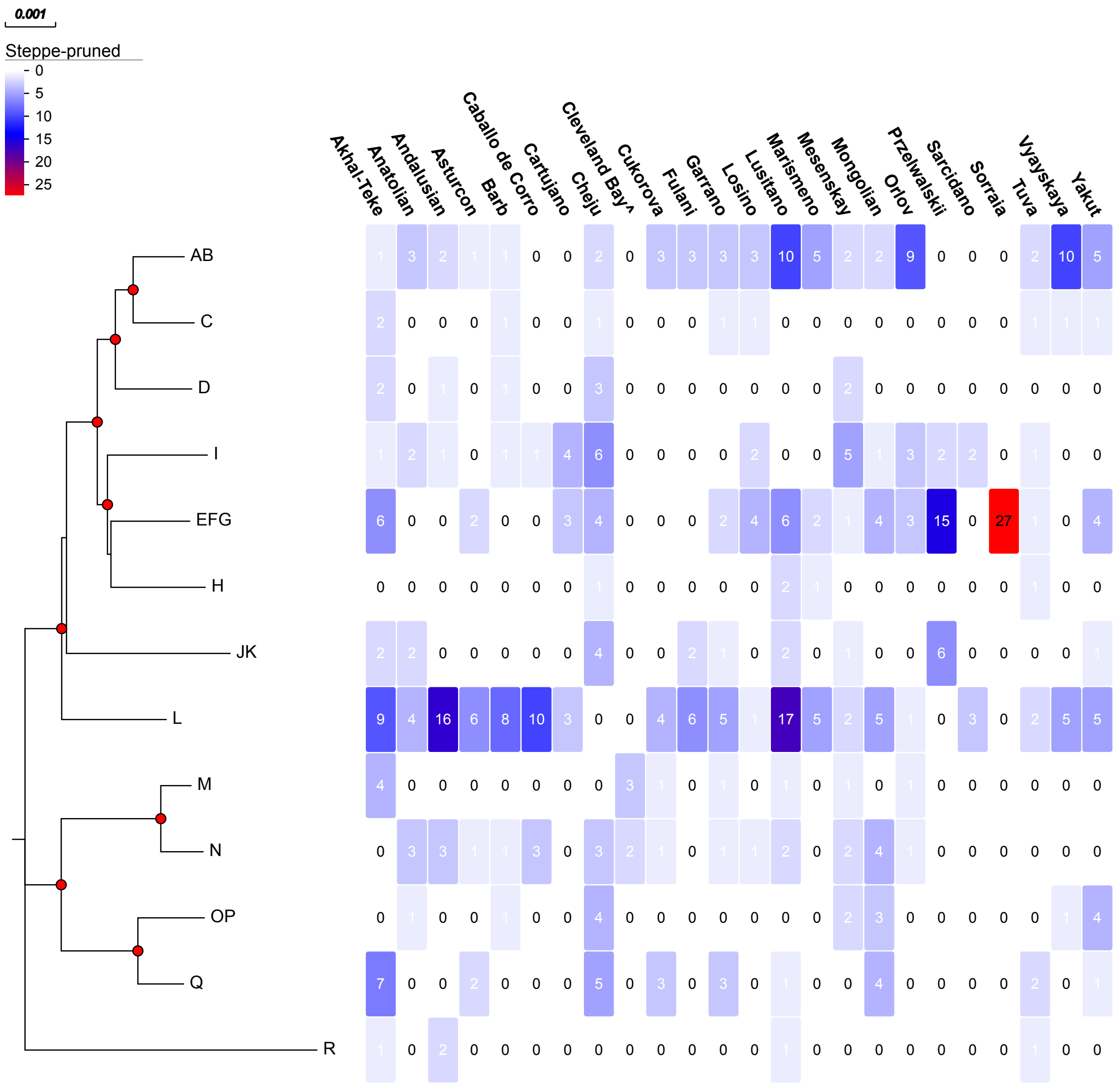


**Supplementary Figure 3. Heatmap showing the relative frequencies of mitogenome haplogroups across breeds classified as Type3: Steppe, visualised using Evolview v2 (He et al. 2016).** The phylogeny on the left shows the haplogroup framework based on a condensed version (one sequence per haplogroup) mitogenome tree (Figure 1); red circles at nodes indicate bootstrap support >90%. Each column on the right is a breed classified as Steppe, in the Southern Group horses (Schafer 1980), showing the number of sequences from each haplogroup. Only breeds with at least 5 sequences available are included. Note that all haplogroups were represented among the Steppe breeds, with the most common being haplogroups L, AB and I. Note that the JK haplogroup that was rare in the northern breeds was found in more of the southern breeds, including Przelwalskii horses. It would be difficult to assign a unique ancestor to breeds morphologically falling into this classification. However, Haplogroup L was lacking in Przelwalskii, Cheju and Cleaveland Bay but haplogroup JK, which is relatively rare among the Steppe breeds, is shared by Przelwalskii and Cheju. Although many breeds likely have mixed origins, indicated are those with suspected influence from: ^ heavy; * steppe; and § proto-Arab horses.

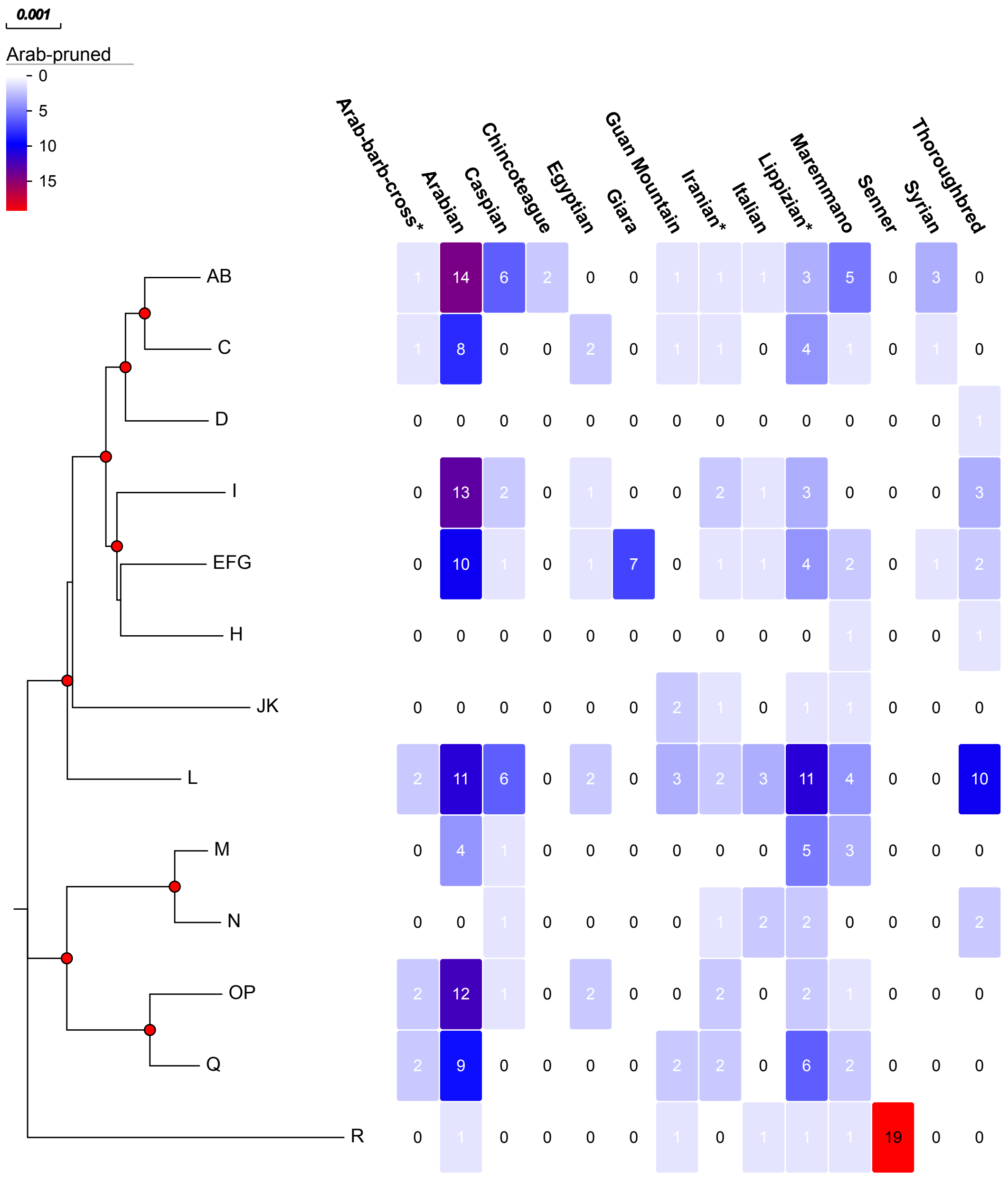


**Supplementary Figure 4. Heatmap showing the relative frequencies of mitogenome haplogroups across breeds classified as Type4: Proto-Arab, visualised using Evolview v2 (He et al. 2016).** The phylogeny on the left shows the haplogroup framework based on a condensed version (one sequence per haplogroup) mitogenome tree (Figure 1); red circles at nodes indicate bootstrap support >90%. Each column on the right is a breed classified as Proto-Arab, in the Southern Group horses (Schafer 1980), showing the number of sequences from each haplogroup. Although haplogroups D and H were rare, all haplogroups were represented among the Proto-Arab breeds, with the most common again being haplogroups L, AB and I. The R haplogroup was found in more of the Proto-Arab compared to the other breeds; all of the Senner sequences sampled shared the same R haplogroup haplotype but it was rare in other breeds. Although many breeds likely have mixed origins, indicated are those with suspected influence from: ^ heavy; * steppe; and § proto-Arab horses.
